# AMPK Phosphorylation Proceeds Through Hierarchical Proteoform Cascades Revealed by Integrated Mass Spectrometry

**DOI:** 10.1101/2025.10.10.681638

**Authors:** Boris Krichel, Hsin-Ju Chan, Liam Bandura, Zhan Gao, Man-Di Wang, Holden T. Rogers, Sean J. Mcilwain, Charlotte Uetrecht, Ying Ge

## Abstract

Protein kinases integrate cellular signals through complex phosphorylation cascades, yet resolving how chemical perturbations trigger and modulate these cascades in therapeutic targets remains a major challenge. Here, we dissect AMP-activated protein kinase (AMPK) proteoforms during activation through controlled biochemical reactions with a hybrid mass spectrometry (MS) approach integrating bottom-up MS for site-specific kinetics with top-down proteoform characterization. We reveal that AMPK phosphorylation proceeds through hierarchical cascades rather than binary switching, with dual entry points: canonical CaMKK2-mediated phosphorylation or allosteric activator PF-739 both triggering extensive autophosphorylation with α1-S496 showing highest kinetic priority. Proteoform-resolved analysis uncovers channeled β1-S24/25+S108 co-phosphorylation linking subcellular localization with allosteric responsiveness. Site-directed mutagenesis demonstrates CaMKK2 targets only α1-T183, with all other modifications arising through autophosphorylation. Phosphatase competition reveals asymmetric control where PP1A selectively removes activation-loop phosphorylation while autophosphorylation sites remain protected, establishing persistent regulatory states. Resolving AMPK’s temporal kinetics and proteoform architecture during activation enables a proteoform-centric understanding on kinase regulation.

## MAIN

Post-translational modifications (PTMs) expand the molecular diversity of the proteome far beyond its genetic blueprint, empowering proteins with the ability to adapt and regulate their functions dynamically^1,2^. Proteoforms, molecular variants arising from PTMs, alternative splicing, and genetic variation, underpin biological complexity^3,4^. Among the PTMs, phosphorylation creates functionally distinct proteoforms that regulate cellular signaling and metabolic homeostasis^5,6^. Elucidating how multiple phosphorylation sites collectively modulate protein function is essential for developing therapeutics targeting metabolic kinases, which control cellular energy homeostasis and represent promising drug targets for metabolic disorders including diabetes and obesity^7,8^. However, capturing the temporal dynamics and combinatorial patterns of these phosphorylations with analytical precision remains challenging^9–11^.

Mass spectrometry (MS)-based proteomics has transformed phosphorylation research by enabling multiplexed and quantitative analysis with sequence-level resolution^12–15^. Bottom-up proteomics excels at deep proteome coverage and site identification through peptide level analysis, yet it disrupts the proteoforms and loses information on combinatorial modification patterns^16,17^. In contrast, top-down proteomics^18^ preserves intact proteoform architectures and combinatorial modification patterns but with limited throughput and difficulty in mapping phosphorylation sites especially for large proteins^19,20^. Resolving phosphorylation cascades demands tracking of how individual sites change over time, because phosphorylation occurs in hierarchical cascades where initial modifications gate subsequent events^21,22^. For kinases with multiple phosphorylation sites and complex regulatory patterns, comprehensive proteoform-centric analysis requires integrating both approaches to capture both the temporal kinetics of individual sites and the combinatorial architecture of intact proteoforms. This integration enables a proteoform-centric perspective on how modification patterns evolve and coordinate regulatory functions, providing understanding inaccessible to single method analyses.

AMP-activated protein kinase (AMPK) is a central regulator of cellular energy metabolism with broad therapeutic relevance across metabolic, cardiovascular, and aging-related diseases^25–28^. This heterotrimeric complex (α-β-γ) is canonically known to require AMP binding and activation loop phosphorylation at α1-T183 by upstream kinases such as calcium/calmodulin dependent protein kinase kinase 2 (CaMKK2) to achieve catalytic activation and downstream substrate phosphorylation^29,30^. However, AMPK also undergoes extensive autophosphorylation, creating diverse proteoform landscapes whose regulatory roles and temporal relationships remain poorly defined^31,32^, exemplifying the challenge of resolving dynamic, multi-site phosphorylation^23,24^.

Here we dissect AMPK’s temporal kinetics and proteoform architecture using an integrated hybrid MS strategy combining quantitative intact mass measurements, bottom-up site-specific kinetics, and top-down proteoform sequencing. By combining kinase phosphorylation, dephosphorylation and allosteric activation with proteoform-resolved MS, we reveal how autophosphorylation creates diverse modification patterns at sites linked to distinct regulatory functions. These findings extend beyond the traditional activation model to define a dynamic and multi-mechanistic proteoform-centric activation model for this major therapeutic target.

## RESULTS

### A hybrid MS strategy resolves proteoform-level phosphorylation dynamics

Our hybrid MS strategy unifies three complementary MS approaches to comprehensively characterize phosphorylation dynamics (Fig. 1a-c); Intact protein reverse phase liquid chromatography mass spectrometry (RPLC-MS) quantifies proteoform changes over time (Fig. 1a), bottom-up MS resolves site-specific phosphorylation kinetics (Fig. 1b), and top-down fragmentation using ultra high-resolution MS definitively identifies proteoforms through sequencing (Fig. 1c). This integrated approach captures temporal kinetics and proteoform architecture by combining complementary mass spectrometry measurements (Fig. 1d).

**Figure 1:**
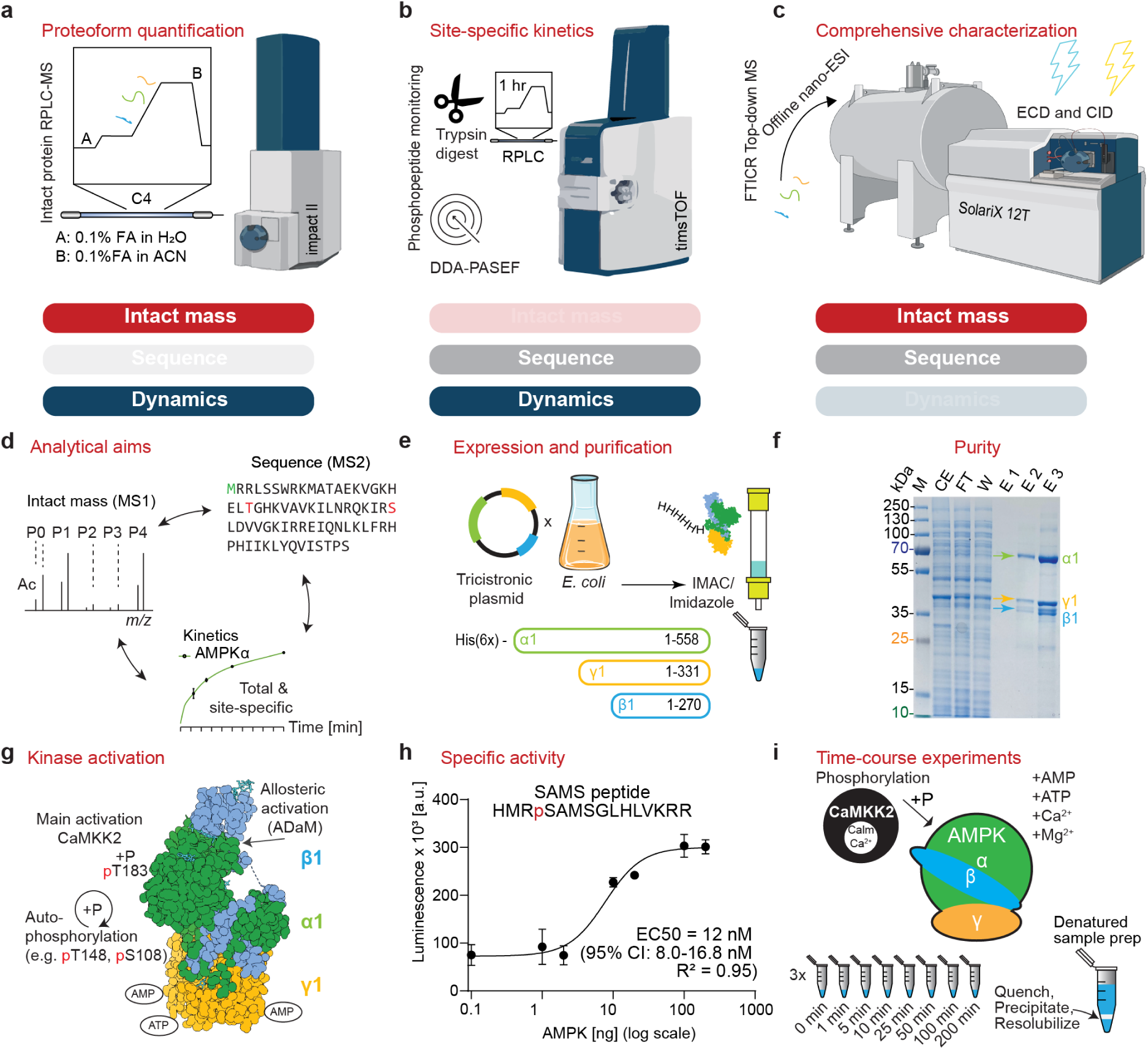
Hybrid mass spectrometry (MS)-strategy for resolving kinase phosphorylation dynamics. **(a)** Intact proteoform tracking via LC-MS. **(b)** Site-specific phosphorylation kinetics via bottom-up MS. **(c)** Top-down proteoform characterization using ultra high-resolution MS with various fragmentation options. **(d)** Conceptual integration of the three approaches to capture intact mass, sequence information, and temporal dynamics. **(e)** Heterologous AMPK expression and affinity purification workflow. **(f)** SDS-PAGE analysis confirming purity of affinity purified AMPK complex with approximately stoichiometric subunit ratios. **(g)** AMPK activation mechanism: AMP binding enhances CaMKK2-mediated T183 phosphorylation and subsequent autophosphorylation cascade. **(h)** Dose-response curve showing AMPK enzyme titration with SAMS peptide substrate. X-axis shows total AMPK amount added to 5 µL reactions on logarithmic scale. **(i)** Time-course phosphorylation workflow: AMPK incubated with CaMKK2 and cofactors, with samples quenched at defined intervals and prepared by chloroform-methanol-water-precipitation.

We applied this strategy to AMP activated protein kinase (AMPK), which exists as 12 possible heterotrimeric combinations through different isoforms (α1-2, β1-2 and γ1-3)^33^. We produced full-length physiological α1β1γ1 isoforms in *E. coli*^34^, the ubiquitously expressed isoform combination in human tissues, and purified the complex to homogeneity (Fig. 1e-f, Table S1). We activated AMPK using AMP and CaMKK2 mediated phosphorylation (Fig. 1g). AMPK functionality was validated through activity assays using synthetic SAMS peptide substrate which confirmed that the enzyme was functional with an EC50 = 12 nM, demonstrating quality comparable to commercial preparations (Fig. 1h)^35^. Phosphorylation reactions were quenched at defined intervals to capture temporal dynamics (Fig. 1i).

### AMPK subunits exhibit distinct phosphorylation profiles

AMPK subunits undergo differential phosphorylation with distinct temporal patterns. To resolve these dynamics, we performed *in vitro* phosphorylation of the AMPK complex using CaMKK2/Calmodulin. CaMKK2 represents a primary upstream kinase of AMPK, phosphorylating α1-T183 in the activation loop, resulting in about 100-fold activity enhancement^29^. To analyze phosphorylation status, we applied an intact protein RPLC-MS approach, providing highly reproducible chromatographic separation (Fig. 2a). Intact mass analysis of unphosphorylated AMPK subunits confirmed the expected molecular weights (Table S2).

**Figure 2:**
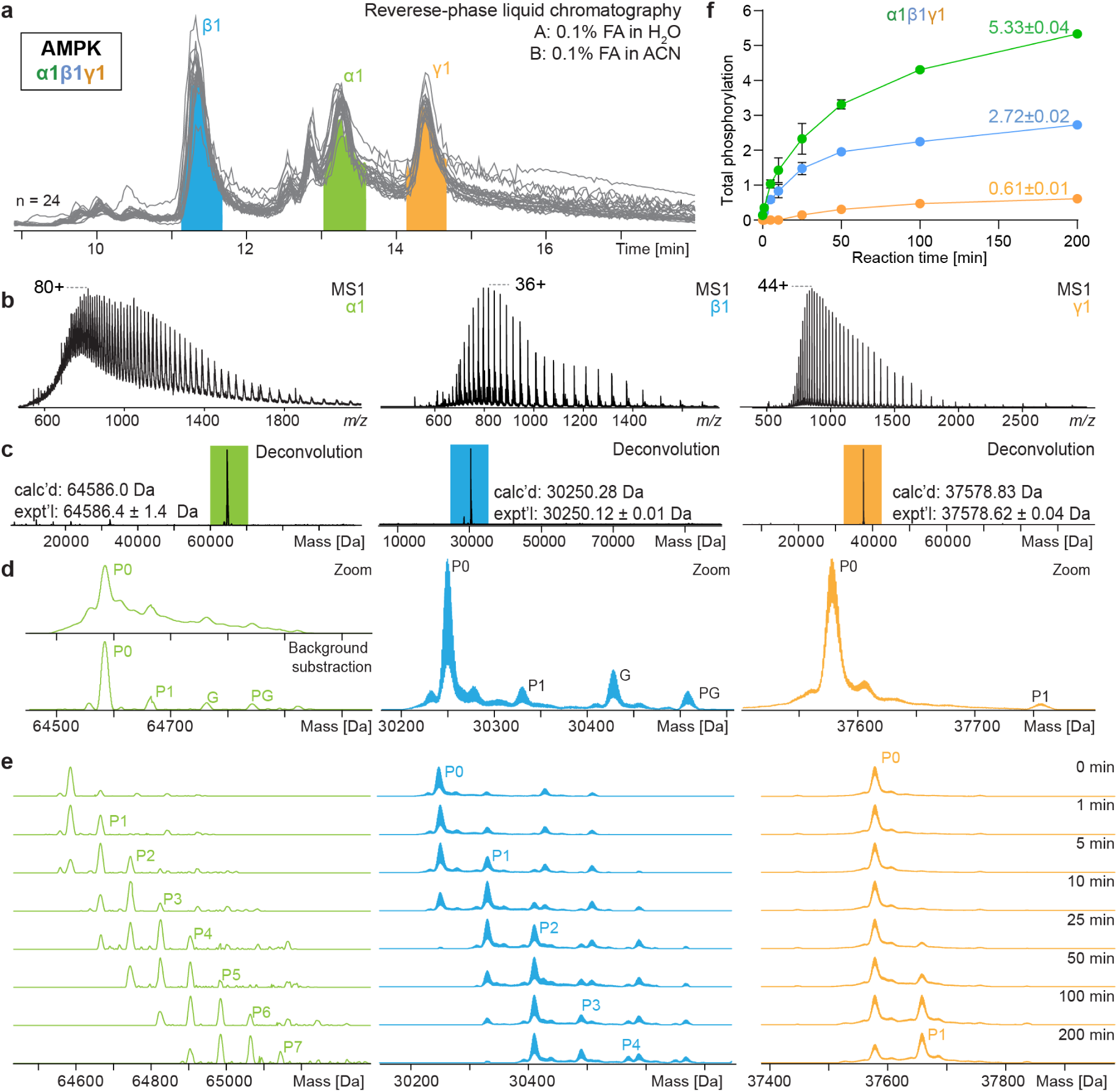
AMPK proteoform quantification by intact protein RPLC-MS during CaMKK2-mediated activation. **(a)** Total ion chromatograms of 24 consecutive runs across the phosphorylation time course. **(b)** MS1 intact protein spectra at 0 min reaction time with most abundant charge states labeled. **(c)** Deconvoluted mass spectra for proteoform identification showing calculated (calc’d) and experimental (expt’l) masses. **(d)** Quantifiable peaks labeled with their corresponding proteoforms. P = Phosphorylation, G = Gluconoylation, PG = Phosphogluconoylation. **(e)** Representative deconvoluted mass spectra showing temporal progression of mass increases corresponding to sequential phosphorylation events. **(f)** Temporal changes in total phosphorylation for each AMPK subunit over 200 min (mean ± SEM, n = 3).

Intact protein MS at the initial time point showed well-resolved charge state distributions (Fig. 2b, Fig. S1a). Deconvoluted mass analysis confirmed proteoform identification with high mass accuracy (Fig. 2c). Although we detected initial phosphorylation and mass heterogeneity through non-specific modifications known from *E. coli*^36^ (Fig. 2d), the unphosphorylated species (0P) was by far the dominant form in all three subunits prior to CaMKK2 treatment, after which phosphorylation levels increased dramatically (Fig. 2e). Analysis of total phosphorylation (P_total_) over 200 min revealed the sequential mass increase across AMPK subunits, culminating in the highest levels for α1 (5.33±0.04), intermediate for β1 (2.72±0.01), and minimal for γ1 (0.61±0.01) (Fig. 2f, extended validation in Fig. S1b).

Representative deconvoluted mass spectra across the time course demonstrate the sequential phosphorylation events (Fig. 2e). The maximum phosphorylation states observed at 200 min (α1: 7P, β1: 4P, γ1: 1P) represent the minimum number of phosphorylatable sites on each subunit, since not all sites reach full stoichiometric phosphorylation. Distribution of AMPK proteoforms throughout the spectra revealed distinct phosphorylation behaviors: α1 exhibited a broad, distributed envelope indicating multiple competing modification pathways, whereas β1 showed preferential accumulation at the P2 state, suggesting a more channeled phosphorylation pattern with one predominant proteoform.

While numerous phosphorylation sites have been reported for AMPK subunits^32,37–41^ and prior intact MS studies have observed activation-dependent proteoform changes in α2β2γ3 complexes^42^, our results reveal richer phosphorylation patterns than expected from direct CaMKK2 targeting alone, indicating triggered autophosphorylation cascades following initial α1-T183 activation. MS-based and biochemical studies report variable numbers of potential autophosphorylation sites (6 for α1, 7 for β1, none for γ1 based on PhosphoSitePlus compilation), although the specific sites and stoichiometry depend heavily on experimental conditions^43^. These quantitative profiles revealed subunit-specific hierarchies, indicating complex regulatory mechanisms beyond canonical T183 phosphorylation.

### Phosphatase competition reveals asymmetric control of AMPK activation

AMPK regulation involves a dynamic equilibrium between kinase-mediated phosphorylation and phosphatase-mediated dephosphorylation, with PP1A preventing constitutive activation^44^. Understanding which sites are preferentially targeted by phosphatases provides insight into AMPK’s regulatory architecture.

We investigated the interplay through competitive phosphorylation-dephosphorylation reactions using increased PP1A concentrations (Fig. 3a). In the absence of PP1A, total phosphorylation after 100 min reached levels (α1: 4.44 ± 0.03P, β1: 2.34 ± 0.01P, γ1: 0.46 ± 0.01P) (Fig. S2a-c) comparable to previous experiments. However, even low PP1A (4.52 µM) concentrations selectively reduced α1 phosphorylation by approximately 1 P (∼0.93 P decrease), while β1 (∼0.18 P decrease) and γ1 levels (decrease within error) remained relatively stable (Fig. 3b). Even at high PP1A concentrations, substantial phosphorylation persisted, indicating that PP1A selectively removes the activating phosphorylation at T183 on the α1 subunit, whereas autophosphorylation sites appear protected from dephosphorylation. This asymmetric control demonstrates that autophosphorylation sites are kinetically stable once established, potentially persisting beyond acute signaling events (Fig. 3c).

**Figure 3:**
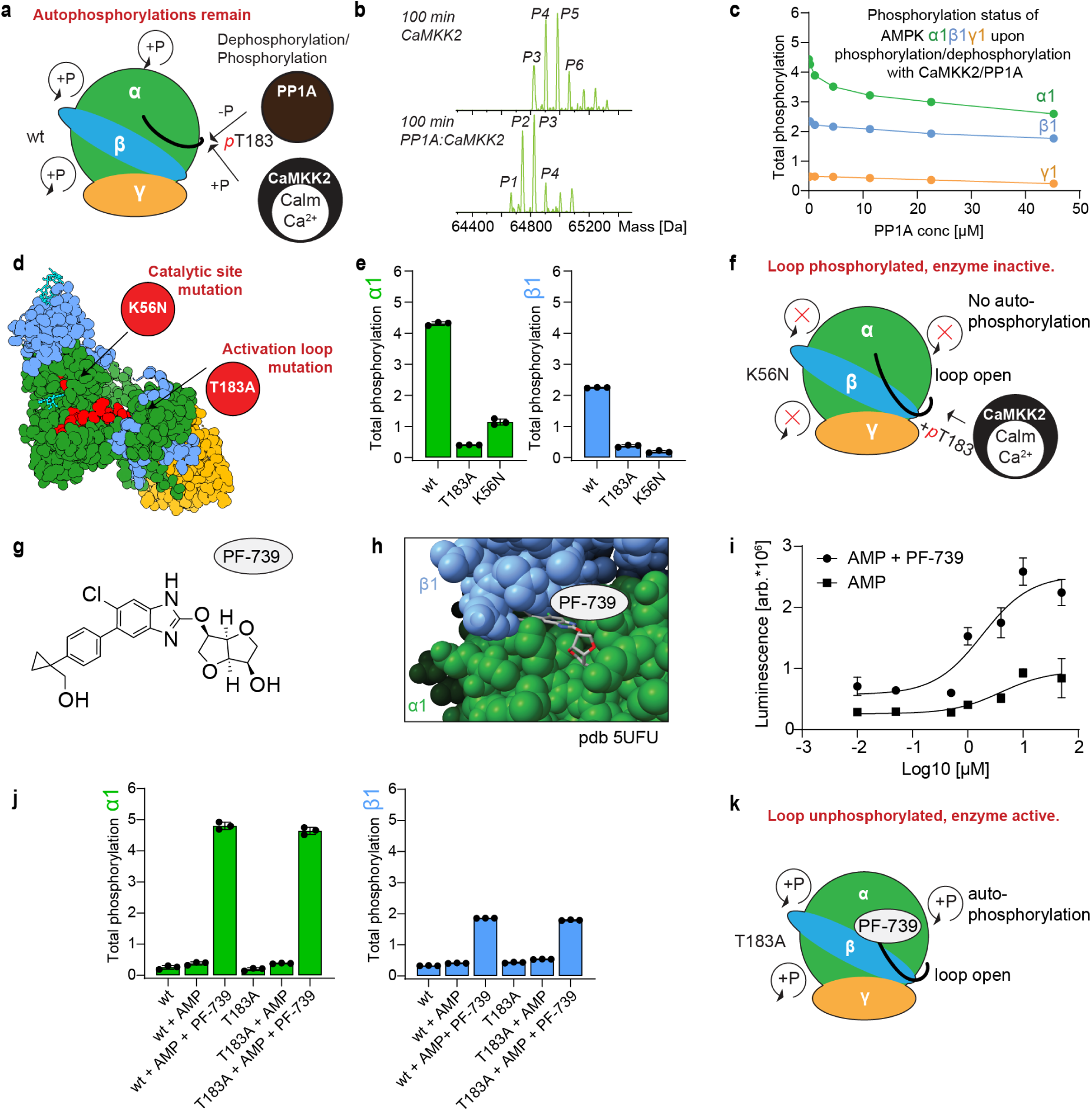
Deciphering canonical and allosteric mechanisms of AMPK activation. **(a)** Schematic showing the competitive phosphorylation-dephosphorylation reactions: CaMKK2-mediated phosphorylation versus PP1A-mediated dephosphorylation of AMPK. **(b)** MS1 intact protein spectra of CaMKK2-phosphorylated AMPK demonstrating that total α1 phosphorylation decreases (∼1P) upon phosphatase (PP1A) addition. **(c)** Concentration-dependent phosphorylation-dephosphorylation dynamics showing that α1 decreases earlier and more substantially than β1 and γ1 subunits (mean ± SEM, n=3). **(d)** Structural model showing site-directed mutations: catalytic site (K56N) and activation loop (T183A). **(e)** Total phosphorylation quantification in WT and mutants following CaMKK2 treatment (mean ± SEM, *n=3*), T183A is inactive upon CaMKK2 addition while K56N increases selectively (∼1 P) in α1 subunit. **(f)** Schematic model illustrating CaMKK2-mediated activation loop phosphorylation as the sole direct target in catalytically inactive K56N mutant. **(g-h)** PF-739 structure and ADaM site binding location. **(i)** ADP-Glo kinase assay showing dose-dependent allosteric activation (arbitrary units, mean ± SEM, *n=3*). Peak activation showing 3.8-fold enhancement over AMP alone. **(j)** Phosphorylation analysis of WT and T183A AMPK following allosteric activation; wt and T183A show intense phosphorylation by allosteric activation, indicating that the T183 phosphorylation is circumvented. **(k)** Proposed mechanistic model illustrating PF-739-mediated enzyme activation despite unphosphorylated activation loop, establishing dual entry into the autophosphorylation cascade.

### Chemical-genetic dissection defines CaMKK2 specificity

To distinguish CaMKK2-driven phosphorylation from AMPK autophosphorylation, we generated two α1-subunit mutants that selectively disrupt different aspects of AMPK activation (Fig. 3d, Fig. S3a). The α1-T183A mutant prevents activation loop phosphorylation, blocking canonical activation. The α1-K56N mutant disrupts ATP binding, abolishing all kinase activity; thus, any phosphorylation detected must originate from an external kinase. This dual-mutant strategy definitively separates upstream kinase targets from downstream autophosphorylation.

RPLC-MS analysis revealed striking differences in CaMKK2-mediated modification patterns (Fig. 3e). Wild-type AMPK exhibited phosphorylation levels (α1: 4.44 ± 0.03P, β1: 2.34 ± 0.01P, γ1: 0.46 ± 0.01P) consistent with previous experiments. The α1-T183A mutant showed virtually no detectable phosphorylation on any subunit, indicating that without T183 phosphorylation, the activation loop cannot open and no autophosphorylation cascade can proceed. Most revealing was the catalytically inactive K56N mutant, which acquired exactly one phosphorylation on α1 and none on β1 (Fig. S3b). This single modification identifies T183 as the sole direct CaMKK2 target. All other phosphorylation sites across AMPK subunits are generated exclusively through autophosphorylation triggered by this initial priming event.

### Allosteric activation bypasses canonical phosphorylation requirements

AMPK harbors an allosteric drug and metabolite (ADaM) binding site at the α-kinase domain and β-subunit interface, activated by small molecules up to 10-fold^45^. We employed PF-739, a chemical probe ADaM site activator, demonstrating dose-dependent AMPK activation with 3.8-fold enhancement over AMP alone (Fig. 3g-i).^46^

We next assessed whether allosteric stimulation alone could initiate the phosphorylation cascade in the absence of the canonical upstream trigger. While AMP alone produced minimal phosphorylation in both wild-type and T183A AMPK, addition of PF-739 resulted in substantial and nearly identical phosphorylation levels in both variants (WT: α1: 4.79 ± 0.12P, β1: 1.79 ± 0.01P; T183A: α1: 4.63 ± 0.12P, β1: 1.79 ± 0.01P) (Fig. 3j, Extended Data Fig. S3c-d). These phosphorylation levels are comparable to those achieved through CaMKK2 activation, demonstrating that allosteric activation enables robust autophosphorylation even when T183 cannot be phosphorylated. This establishes that activation loop phosphorylation is not strictly required for AMPK to execute its regulatory phosphorylation program. These results support a model where PF-739 binding promotes a catalytically competent conformation that bypasses the canonical requirement for activation loop phosphorylation, establishing dual entry points into the hierarchical autophosphorylation cascade (Fig. 3k, Fig. S4a)^47^.

### Site-specific phosphorylation kinetics reveal mechanistic hierarchy

To map the kinetic hierarchy underlying AMPK proteoform dynamics, we employed bottom-up MS using identical aliquots from the CaMKK2-AMPK reaction samples (Fig. 4a, Fig. S5b-d). Phosphorylation kinetics were quantified by monitoring extracted ion chromatograms (EIC) for peptide pairs showing inversely correlating behavior (Fig. 4b-c, Table S3-5, Fig. S5a). Monitoring unphosphorylated peptide disappearance for sixteen peptide pairs provided the most reliable measure of modification progression (Fig. 4d).

**Figure 4:**
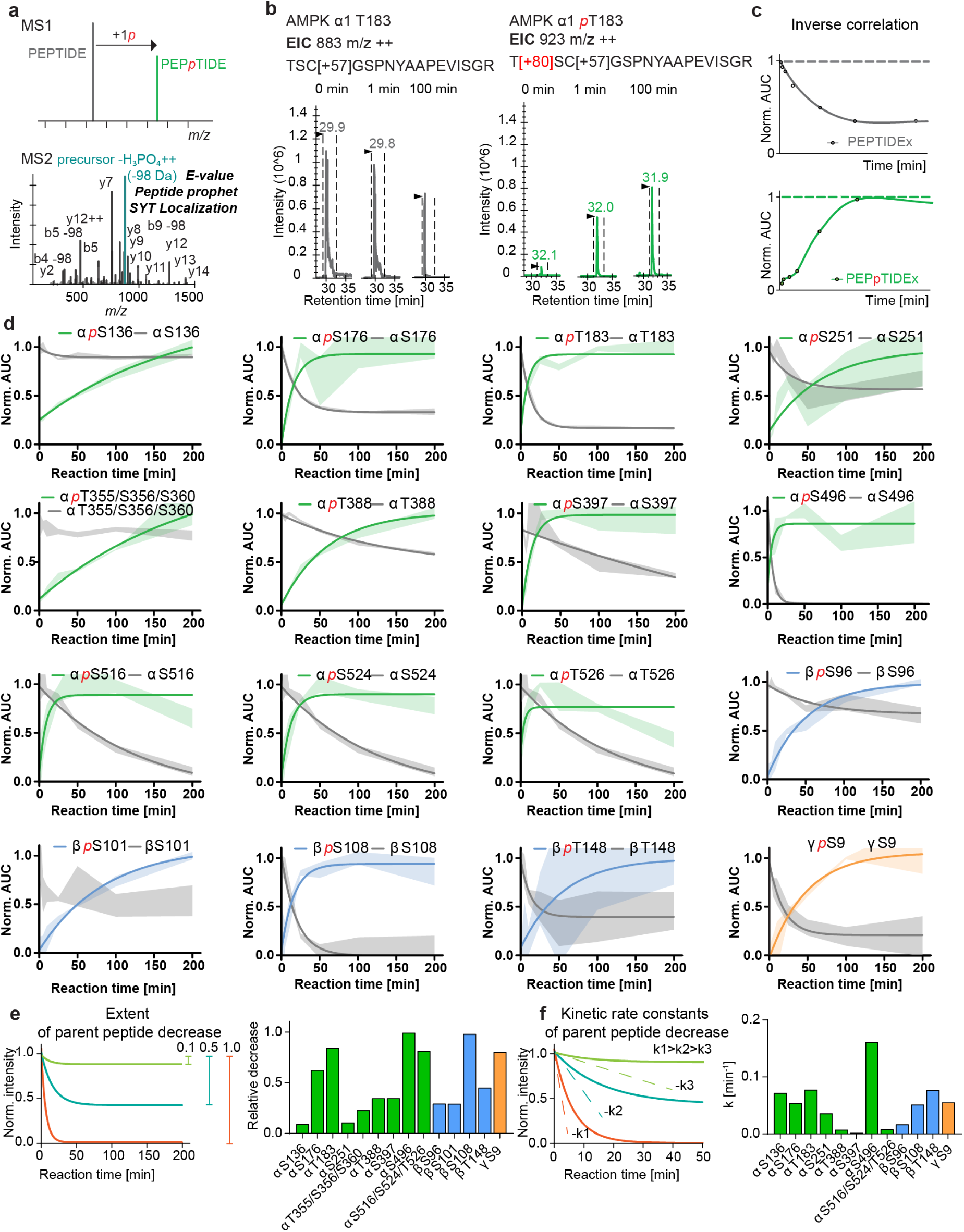
Site-specific phosphorylation kinetics reveal mechanistic hierarchy using bottom-up MS. **(a)** MS1 **of** phosphorylated/unphosphorylated peptide pair tracking and MS2 spectrum showing spectral quality and phosphopeptide identification. **(b)** Temporal progression at 0, 1, and 100 min for parent peptide and phosphopeptide T183 showing inversely correlating intensities. **(c)** Schematic of kinetic plots with inversely correlating changes over time monitored as Normalized Area Under Curve (AUC). **(d)** Site-specific phosphorylation kinetics for 16 peptide pairs with color-coded subunit assignment (α1: green, β1: blue, γ1: yellow-orange). One-phase decay curve fitting applied. Shaded bands represent mean ± SEM. **Note**: S516, S524 and T526 have the same parent peptide. Parent peptides of α1 T355/T356/S360 and β1 S101 could not be fit with the one-phase decay model. **(e)** Schematic showing how extents of phosphorylation were determined from parent peptide decrease; bar graph showing extents of phosphorylation. **(f)** Schematic showing how kinetic rate constants (k) were obtained by fitting peptide disappearance curves to a one-phase decay model. **Note**: Parent peptides for β1 S101 and α1 S397 showed early-timepoint variability within the first 10 min, could not be fit to the one-phase decay model and were only analyzed for their extent. The α1 S397 quantification challenges likely arise from (1) the peptide containing one missed cleavage site, complicating accurate quantification, and (2) sequence overlap with the T388-containing peptide, where T388 phosphorylation also decreases the S397 parent peptide intensity.

Kinetic profiling revealed a distinct hierarchy of phosphorylation rates across AMPK sites (Fig. 4e-f). The most rapid autophosphorylation occurred at α1-S496, an inhibitory site^48^, exhibiting exceptionally high kinetic rate (0.16 ± 0.05 min⁻¹). This reflects efficient trans-autophosphorylation where CaMKK2-activated AMPK molecules phosphorylate S496 on neighboring complexes faster than CaMKK2 phosphorylates α1-T183 (0.077 ± 0.005 min⁻¹).

The well-characterized β1-S108 showed excellent kinetic behavior (R² = 0.99) with complete modification by 50 minutes^47^. β1-T148, crucial for glycogen binding^49^, exhibited higher rate constant (0.077 ± 0.083 min⁻¹) but moderate extent (0.45) and quantification challenges from complex peptide sequence, making kinetic parameter interpretation less reliable^49^. These kinetic profiles established a temporal hierarchy distinguishing rapid regulatory modifications from slower secondary events.

Several sites displayed moderate kinetics, including α1-S176 (0.053 ± 0.014 min⁻¹, extent = 0.62) and γ1-S9 (0.055 ± 0.027 min⁻¹, extent = 0.80), while sites like α1-T388 (0.007 ± 0.004 min⁻¹, extent = 0.35) showed very slow kinetics likely representing secondary regulatory mechanisms.

Among the analytical challenges was quantification of convoluted phosphorylations at the α1 C-terminal regulatory domain. However, a shorter 23-residue peptide from semi-specific trypsin cleavage (SSEVSLTSSVTSLDSSPVDLTPR/P) separated into three distinct chromatographic peaks corresponding to phosphorylation at α1-S516, α1-S524, and α1-T526 (Fig. S5e). The unphosphorylated peptide decreased slowly (k = 0.008 ± 0.008 min⁻¹) but achieved high phosphorylation extent (0.81). The total phosphorylation is distributed across three sites with chromatographic peak area ratios (S516 : S524 : T526 = 1.0 : 1.9 : 0.7) indicating preferential S524 modification.

The α1 and β1 subunit difference between peptide-level and intact protein quantification (4.6P vs. 5.3P, 2.0P vs. 2.7P, respectively) suggests the presence of modifications not captured by bottom-up analysis. Site-specific kinetic profiling revealed AMPK autophosphorylation proceeds through a hierarchical cascade with distinct temporal classes. While this identified individual site kinetics, the specific combinations of co-occurring modifications within intact proteoforms remained unresolved, motivating direct proteoform characterization.

### Proteoform-resolved characterization identifies channeled modification patterns

To identify co-occurring modifications and resolve proteoform architectures, we employed ultrahigh resolution (>400,000 at 400 *m/z*) MS top-down sequencing analysis. Direct Nano-ESI of denatured AMPK subunits yielded high-quality spectra with isotopic resolution. However, the α1 subunit showed strong signal suppression due to size-related effects and was only detectable in the unphosphorylated control sample. In contrast, the smaller β1 and γ1 subunits dominated the signal response in both conditions (Fig. 5a, Fig. S6a-b).

**Figure 5:**
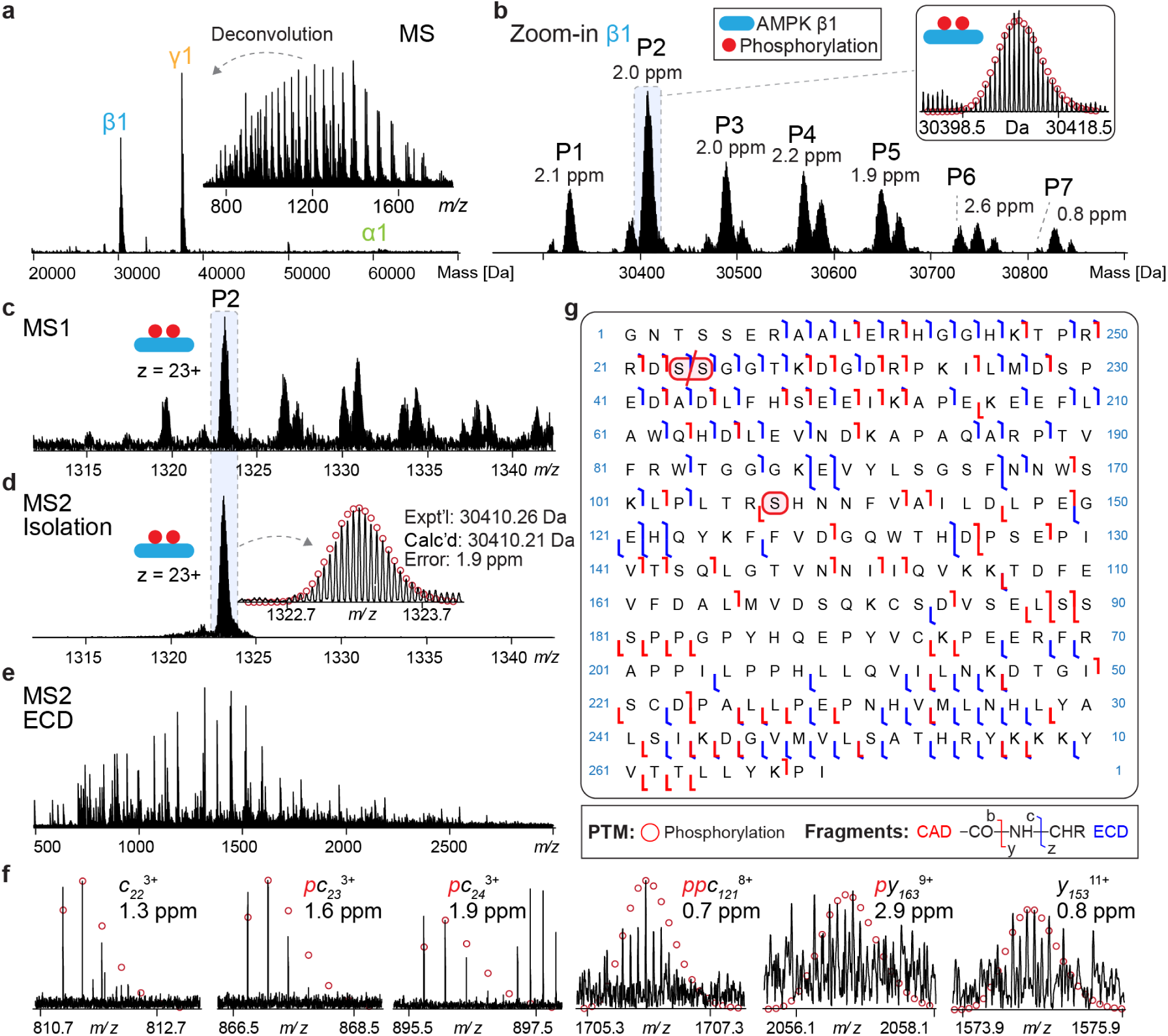
Top-down sequencing identifies doubly phosphorylated AMPK β1 proteoform architecture. **(a)** Deconvoluted mass spectrum of denatured AMPK subunits with raw mass spectrum shown in inset. **(b)** Zoom-in shows isotopically resolved β1 phosphorylation states (1P-7P) with 2P as the most abundant species. **(c)** The 23+ charge state of the β1 shows a peak distribution similar to the deconvoluted spectrum in (b). **(d)** Quadrupole isolation of 2P β1 23+ precursor ion (1323 *m/z*) with isotopically resolved peak inset. **(e)** Representative ECD fragmentation spectrum showing extensive fragmentation pattern. **(f)** Selected fragment spectra (*p*: singly phosphorylated; *pp*: doubly phosphorylated). Mass errors are reported, and isotopic fittings are shown in red circles. Shift from fragment pattern *c_22_, c_23_, c_24_* versus assigned positions *p*S24, *p*S25 due to N-terminal glycine cleavage. **(g)** Combined fragment map from complementary CID and ECD experiments. Fragment notation: hook direction indicates type, color denotes method (blue: ECD, red: CID).

Examination of β1 intact mass spectra from the CaMKK2-treated sample revealed heterogeneous phosphorylated species (1P-7P), with doubly phosphorylated (2P) as the major component (Fig. 5b). Top-down measurements showed total β1 phosphorylation of 3.1P, slightly higher than our earlier RPLC-TOF analysis (2.7P), likely due to different ionization conditions. Summing phosphorylation extents from all identified peptides in bottom-up analysis yielded only 2.0P, revealing a 0.7-1.1P discrepancy compared to intact protein measurements. This discrepancy made the predominant 2P β1 proteoform an ideal target for identifying the missing modifications through direct sequencing. Quadrupole isolation of the 2P β1 23+ precursor (1323 *m/z*) achieved mass species selectivity with high mass accuracy (1.9 ppm) (Fig. 5c-d).

Comprehensive structural characterization was performed using complementary fragmentation approaches combining collision-induced dissociation (CID) and electron capture dissociation (ECD) at optimized parameters (Fig. 5e, Fig. S7). Data processing used in-house MASH software with manual curation.^50^ The combined fragmentation experiments yielded 179 high-confidence fragment ions (43 *b-*, 54 *c-*, 34 *y-*, 48 *z-*type) across the 269-residue β1 sequence, achieving 54% backbone coverage and high mass precision (Figure 5f-g). Systematic fragment analysis revealed the major β1 bis-phosphorylated proteoform (2P) carries phosphorylation at S24/25 and S108.

The first phosphorylation was localized to the S24/25 double serine motif based on c22, c23, and c24 fragment ions, with predominant modification at S24 (Fig. S8), consistent with the mutually exclusive phosphorylation pattern described for this site.^40^ The identification of S24/25 phosphorylation directly addresses the quantitative discrepancy between intact protein and peptide-level measurements. This site was absent from bottom-up analysis due to poor N-terminal coverage; the 6-residue tryptic peptide DSSGTK containing S24/S25 was not detected.

The second phosphorylation site, S108, was definitively assigned based on 27 high-confidence fragment ions spanning this region. This S24/25+S108 combination represents a notable finding, as S24/25 phosphorylation has been linked to extranuclear distribution, while S108 phosphorylation facilitates allosteric activation by stabilizing the ADaM site binding pocket^38,51^. The strong predominance of the 2P species in intact mass spectra, combined with the consistent identification of this specific S24/25+S108 combination, suggests a channeled proteoform formation pathway rather than stochastic modification of available sites. In contrast, β1-S101 and β1-T148 phosphorylation could not be detected in our top-down analysis, despite identification in bottom-up experiments, indicating these represent minor proteoforms under these experimental conditions. Proteoform-resolved characterization thus revealed selective modification patterns that coordinate distinct regulatory mechanisms within AMPK.

### Integrated analysis reveals mechanistic model of AMPK activation

Integrating kinetic, proteoform, and chemical perturbation data reveals a model for AMPK proteoform regulation based on four key mechanisms (Fig. 6a): (1) Dual entry into the activation cascade through CaMKK2-dependent α1-T183 phosphorylation or allosteric activation. (2) Hierarchical phosphorylation kinetics with defined priorities, where α1-S496 shows highest modification efficiency. (3) Channeled proteoforms with selective co-occurrence, exemplified by the predominant β1-S24/25 + S108 combination coupling localization and allosteric sensitivity. (4) Asymmetric dephosphorylation control where PP1A reverses α1-T183 but not autophosphorylation sites, enabling proteoform persistence beyond acute signaling.

**Figure 6.**
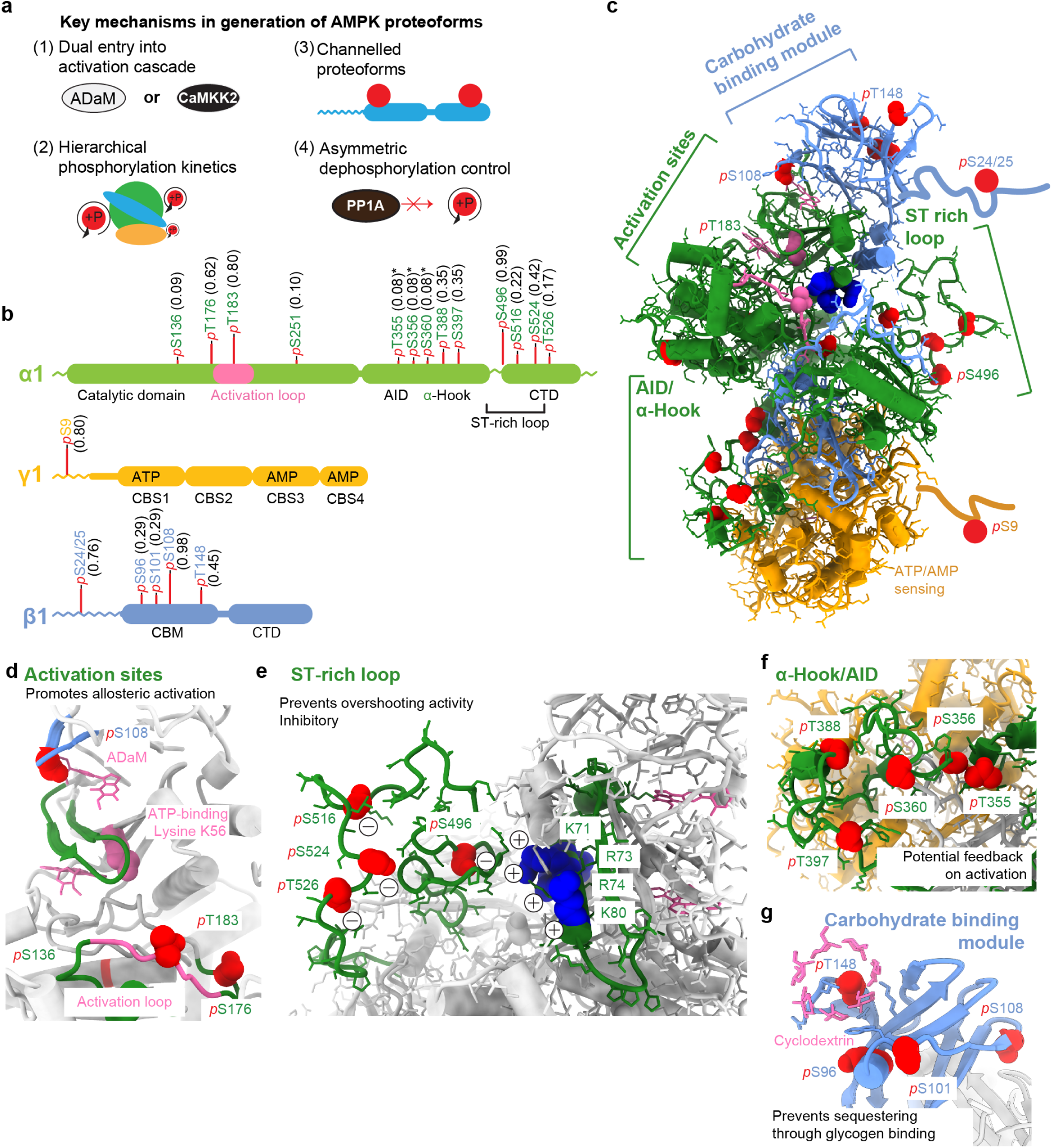
Spatial organization and regulatory mechanisms of AMPK phosphorylation sites revealed by integrated proteoform analysis. **(a)** Summary of mechanisms contributing to AMPK proteoform diversity. **(b)** Domain organization of AMPK α1β1γ1 subunit isoforms showing main functional domains and positions of identified phosphorylation sites. Red bar heights reflect relative extent of phosphorylation determined from kinetic analysis. Four CBS domains with nucleotide binding pockets here depicted in occupational state of activated enzyme. C-terminal domain (CTD). *Asterisked sites were co-identified on the same peptide and assigned equal phosphorylation stoichiometry. **(c)** AMPK α1β1γ1 protein structure homology model (Swiss-Model) based on PDB 5EZV and 4RER. Disordered termini represented as lines. Red spheres indicate identified phosphorylation sites clustered into four distinct structural hotspots. **(d)** Activation sites and catalytic center hotspot. Phosphorylations proximal to activation sites have critical regulatory functions. **(e)** ST-rich loop hotspot. Phosphorylation sites within this regulatory disordered region create electrostatic interactions with basic residues near the catalytic center, reducing T183 phosphorylation by upstream kinases and preventing activity overshooting through negative feedback. **(f)** Autoinhibitory domain (AID) and α-hook hotspot. This region facilitates allosteric communication between AMP binding at the γ-subunit and kinase domain activation. The identified phosphorylation cluster (T355, S356, S360, T388, S397) may constitute a feedback mechanism modulating AMP sensitivity and allosteric regulation. **(g)** Carbohydrate-binding module (CBM) hotspot. The CBM mediates AMPK sequestration at glycogen matrices. While β1-T148 has been functionally characterized for glycogen binding regulation, *p*S96 and *p*S101 remain uncharacterized but may influence CBM function due to structural proximity.

Phosphorylation sites cluster into four structural hotspots with distinct regulatory functions (Fig. 6b-c). The activation sites and catalytic center (Fig. 6d) cluster α1-T183, α1-S176, and β1-S108, where α1-T183 and β1-S108 drive main and allosteric activation. The α1 ST-loop (Fig. 6e) encompasses S496, S524, S516, and T526, orchestrating negative feedback through electrostatic interactions^48,52^. The α1-auto-inhibitory domain and α-hook region (Fig. 6f) contain phosphorylations at T388, S397, S360, S356, and T355^53^. The β1 carbohydrate-binding module (Fig. 6g) features T148 mediating glycogen sequestration^49^, alongside S96 and S101. This spatial organization reveals how AMPK integrates multiple regulatory inputs through autophosphorylation cascades that create proteoform diversity, extending beyond a binary activation model to enable fine-tuned kinase activity and function.

## DISCUSSION

Protein kinases regulate cellular processes through dynamic phosphorylation cascades, yet understanding how chemical perturbations trigger and control these cascades requires resolving both temporal kinetics and proteoform architecture. Applied to AMPK, our hybrid mass spectrometry approach combining intact protein profiling, site-specific kinetics, and proteoform characterization reveals when sites are modified, how modifications propagate hierarchically, and which combinations form co-occurring proteoforms^23,29^.

We discover that AMPK activation proceeds through hierarchical autophosphorylation cascades triggered by a single priming event. Kinetic profiling resolved distinct temporal classes: rapid modification of α1-S496, moderate rates at allosteric control sites^32^, and slow modifications clustered in regulatory domains. These findings reveal for the first time the hierarchical ordering during activation. Phosphatase experiments revealed asymmetric regulation where PP1A selectively removes activation-loop phosphorylation while leaving autophosphorylation sites intact. Prior studies observed one autophosphorylated site corresponding to α1-S496 being resistant to protein phosphatases using western blot^40^, yet, our intact protein measurements demonstrate that specific deactivation does not roll back the proteoform diversity generated through activation.

Allosteric activation revealed a second activation route. PF-739 initiated robust autophosphorylation without T183 phosphorylation. Although allosteric activation without T183 phosphorylation may have limited physiological relevance for canonical AMPK function^47^, the substantial proteoform remodeling suggests diverse modification states can be generated through alternative pathways with implications for selective pharmacological modulation. Proteoform characterization identified channeled β1-S24/25+S108 co-phosphorylation as the predominant modified state, coupling subcellular localization with allosteric responsiveness through selective site co-occurrence^38,49^.

Integrating complementary measurements revealed modifications missed by individual approaches. Sites missed in bottom-up workflows were recovered by intact and top-down analysis, while discrepancies exposed biases obscuring regulatory logic. However, our in vitro measurements do not capture cellular context: in vivo proteoform distributions will depend on cell type, subcellular localization, metabolic state, and local kinase and phosphatase concentrations^24^.

In summary, our findings establish that AMPK operates through hierarchical proteoform cascades that enable fine-tuned regulation beyond binary switching. Combining controlled biochemical reactions with hybrid MS to capture temporal kinetics, proteoform architecture, and responses to chemical and genetic perturbations revealed regulatory mechanisms inaccessible to any single method. Dual activation pathways, hierarchical autophosphorylation kinetics, and selective proteoform co-occurrence identify nodes for pharmacological intervention. These discoveries demonstrate that resolving the kinetics and architecture of proteoform cascades is essential for dissecting complex regulatory mechanisms in AMPK and other therapeutic kinase targets in metabolic disease.

## METHODS

### Genes and Protein Constructs

Recombinant protein sequences were based on UniProt entries human AMPK alpha1 (Q13131), beta1 (Q9Y478), and gamma1 (P54619). The expression plasmid pET28-H6-AMPKγ1β1α1 for AMPK full-length, with His tag (MSYYHHHHHHEGVRTM-) N-terminally fused to the alpha subunit, and pET24 H6GST-hPP1Aα (aa1-314, PP1A domain) was generated by Karsten Melcher Lab and kindly provided by Van Andel Institute. Site-directed mutations on AMPK α K56N and T183A were introduced using the Q5 site-directed mutagenesis kit (New England Biolabs) and confirmed using long-read DNA sequencing (UW-Madison Biotechnology Center).

### Expression and Purification

AMPK and PP1Aα were produced as recombinant proteins in One Shot BL21 Star (DE3) E. coli (Thermo Fisher). Cells were grown in 1 L baffled flasks to OD₆₀₀ = 0.8-1.0, induced with 0.1 M Isopropyl β-D-1-thiogalactopyranoside and continued at 18 °C for 20 h. Cell pellets were lysed in His equilibration buffer (20 mM imidazole, 40 mM phosphate, 300 mM NaCl, pH 8) by sonication. His-tagged proteins were purified using HisPur Ni-NTA resin (Thermo Fisher) with sequential washes (50 mM imidazole) and elution (300 mM imidazole). Proteins were buffer-exchanged into storage buffer (20 mM phosphate, 150 mM NaCl, 300 mM imidazole, 15% glycerol, 2 mM DTT), concentrated to 5-10 mg/mL, and stored at -80 °C. Concentrations were determined by Bradford assay (BioRad) or NanoDrop (Thermo Scientific). CaMKK2 (CaMKK beta isoform 2 [1 – 541]) was obtained through the MRC PPU Reagents and Services facility (MRC PPU, College of Life Sciences, University of Dundee, Scotland):

### AMPK Phosphorylation Reactions

All reactions were performed in assay buffer (50 mM HEPES, 100 mM NaCl, 2 mM DTT, 0.01% Brij 35). CaMKK2 was premixed with calmodulin (Enzo Life Sciences) at a 1:2 molar ratio. For phosphorylation, 7.5 μM AMPK was incubated with 0.155 μM CaMKK2 and 0.31 μM calmodulin in the presence of 500 μM ATP, 200 μM AMP, 5 mM (CH₃COO)₂Mg, and 1 mM CaCl₂ at 20 °C for up to 200 min in [240 μL total volume]. Aliquots (∼15 μg protein) were quenched at defined intervals with 0.1% formic acid (FA). For concentration-dependent PP1A phosphatase experiments, PP1A was added at varying concentrations to CaMKK2 AMPK mixtures, and samples were quenched after 200 min.

Quenched reactions were desalted using chloroform-methanol-water precipitation. AMPK was diluted into 400 μL water, precipitated with 400 μL methanol, phase-separated with 100 μL chloroform, and centrifuged at 15,000 g for 10 min. Following supernatant removal, 400 μL methanol was added and mixed by rocking. After centrifugation, proteins were further desalted by two consecutive methanol washes. Air-dried pellets were stored at -80°C until analysis.

### Intact LC-MS Analysis

Desalted reaction samples were reconstituted in 5% FA/20% acetonitrile-isopropanol and adjusted to 200 ng/μL. LC-MS was performed using nanoAcquity UPLC (Waters) coupled to Impact II Quadrupole Time-Of-Flight (QTOF) (Bruker Daltonics). Samples (300-500 ng) were separated on a home-packed C4 column (Halo, 2.7-μm, 250-μm ID, 10 cm, 1,000 Å) using gradient: 0-3 min, 10% B; 3-20 min, 10-80% B; 20-24 min, 80-95% B; 24-30 min, 95%-10% B (A: 0.1% FA in H₂O; B: 0.1% FA in acetonitrile) at 4 μL/min.

Mass spectra were acquired at 0.5 Hz over 300-3000 m/z. Source parameters: capillary voltage 4500 V, nebulizer pressure 0.5 bar, dry gas 4.0 L/min at 200°C. Ion transfer settings: funnel RF 300 V, quadrupole energy 5 V, low mass cutoff 650 m/z, isCID 10 V, collision RF 2600 Vpp.

Data processing used Compass Data Analysis (Bruker). Summed mass spectra were generated from fixed time windows. Deconvolution employed MaxEnt with different parameters for isotopic resolved (β/γ: resolution 40,000) versus non-resolved (α: resolution 10,000) charge envelopes. Spectra were smoothed (Savitzky-Golay 0.2), background subtracted (0.55), and peak-picked. Peak lists were manually curated for phosphorylation masses and exported for weighted average calculations. The total weighted phosphorylation average was calculated by the following formula:

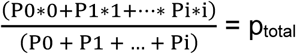

### Bottom-up MS Analysis

Desalted reaction samples were reconstituted and neutralized with ammonium bicarbonate (pH 8), before being reduced with 10 mM TCEP (37°C, 30 min), alkylated with 500 mM chloroacetamide (RT, 30 min), and digested with trypsin (37°C, overnight). Peptides were desalted using C18 stage tips and analyzed by LC-MS/MS.

Samples (200 ng) were separated on Ion Optics columns (25 cm × 75 μm, C18 1.6 μm) at 55°C using 50 min gradients (2-37% B, then 37-85% B over 5 min; A: 0.2% FA in water, B: 0.2% FA in ACN) at 400 nL/min. Analysis used a Trapped Ion Mobility Spectrometry (tims)-TOF Pro (Bruker) in PASEF mode with ion mobility 0.60-1.60 Vs/cm² and standard fragmentation parameters (10 MS/MS scans, 1.17 s cycle time, 20-59 eV collision energy).

Data were processed in Skyline (v23.1.0.380) to extract peak areas (area under the curve, AUC) from corresponding phosphorylated/unphosphorylated peptide pairs. Peak areas were normalized to the maximum value within each peptide series and analyzed in GraphPad Prism. Unphosphorylated peptides were fit to one-phase decay models to obtain rate constants (k, min⁻¹), while phosphorylated peptides were fit to one-phase association models for validation. Phosphorylation extent was calculated as (AUC_max_ - AUC_min_)/AUC_max_ from the normalized peak areas, representing the fraction of peptide phosphorylated. Only peptide pairs showing inverse correlation were included in the final analysis.

Phosphorylation kinetics were determined from unphosphorylated peptide disappearance (normalized to initial values), as this provides the most reliable measure of modification progression independent of differential ionization efficiencies between modified and unmodified forms.

### Top-down MS

Phosphorylated AMPK samples were analyzed using solariX XR 12T Fourier-Transform Ion Cyclotron Resonance (FTICR) mass spectrometer (Bruker) coupled to TriVersa Nanomate (Advion). NanoESI conditions: 0.45 PSI desolvating gas pressure, 1.5-1.55 kV voltage, 3 L/min dry gas at 180 °C. Source optics were optimized: capillary exit 240 V, detector plate 220 V, funnel 1 at 150 V, skimmer 50 V, funnel RF 300 Vpp, octopole frequency 5 MHz with 600 Vpp RF, collision cell at 2 MHz with 2000 Vpp RF.

Mass spectra were acquired with 2M data points over 200-4000 m/z range. Precursor isolation used ≥4 m/z windows. Collision-activated dissociation (CAD) employed 12-35 V collision energy. Electron capture dissociation (ECD) parameters: pulse length 0.01-0.03 s, bias 0.5-1.5 V, lens voltage 5-15 V. Fragment ion assignments were performed using standard top-down proteomics approaches for proteoform characterization. MS2 data were analyzed using MASH Native for proteoform sequencing and PTM localization^50^. Fragments were identified using eTHRASH algorithm. All fragments were manually validated.

### Data Integration and Analysis

Intact protein phosphorylation levels were quantified from deconvoluted mass spectra using weighted averages. Site-specific extent from bottom-up analysis were correlated with total phosphorylation measurements. FTICR MS2 data provided definitive proteoform identification and phosphosite localization and used for complementing the results.

## DATA AVAILABILITY

The mass spectrometry proteomics data generated in this study have been deposited to the ProteomeXchange Consortium via the MassIVE repository with identifier MSV000099446.

## ACKNOWLEDGEMENTS

B.K. was supported by the European Union grant 101068151, Top-AMPK, HORIZON-MSCA-2021-PF-01. B.K. thanks Mel Park for co-supervising the Top-AMPK project. H.T.R. would like to acknowledge support from the National Heart, Lung, and Blood Institute of the NIH under Award Number T32HL007936 through the UW-Madison Cardiovascular Research Center and the Ruth L. Kirschstein NRSA Individual Predoctoral Fellowship under Award Number F31HL178305. C.U. would like to acknowledge funding from the ARIADNE VIBE project, Grant Agreement No. 964553. Y.G. would like to acknowledge NIH R01HL109810, R01GM117058, and S10 OD018475 (to Y.G). The author(s) thank the University of Wisconsin Carbone Cancer Center Cancer Informatics Shared Resource, supported by P30 CA014520 for use of its services. All mass spectrometry data reported in this study were acquired at the University of Wisconsin–Madison Human Proteomics Program Mass Spectrometry Facility. We thank Dr. Yanlong Zhu, the facility manager, for expert instrument support and assistance.

## AUTHOR CONTRIBUTIONS

Y.G. conceived the study. B.K., C.U., and Y.G. designed the experiments; Y.G. supervised the experimental work; B.K., L.B. expressed and purified proteins; B.K. and H-J.C. established and performed phosphorylation assays and MS sample preparation under Y.G.’s direction; H-J.C. acquired FTICR top-down MS fragmentation data; B.K., H.T.R and Z.G. acquired Impact II top-down MS intact data; B.K. and H-J.C. acquired timsTOF bottom-up data; B.K. and H-J.C. analyzed and visualized MS data; B.K. and M.W. performed kinase activity assays; B.K. and S.J.M. calculated total phosphorylation and charge state distribution. Y.G. and C.U. supervised the project; B.K., H-J.C, C.U. and Y.G. evaluated results; B.K., H-J.C, C.U. and Y.G. wrote the paper with comments from other co-authors.

## CONFLICT OF INTEREST

The authors declare no competing financial interests or conflicts of interest in relation to the work described in this manuscript.

## Notes

### Competing Interest Statement

The authors have declared no competing interest.

### Summary of Updates

Revised manuscript for journal submission. Key changes: (1) Updated title to emphasize AMPK findings, (2) Reframed abstract and introduction to highlight chemical perturbations and therapeutic relevance, (3) Streamlined discussion to focus on AMPK regulatory mechanisms, (4) Minor editorial improvements throughout.

