## SUPPLEMENTAL INFORMATION for "AMPK Phosphorylation Proceeds Through Hierarchical Proteoform Cascades Revealed by Integrated Mass Spectrometry"

### Supporting Information for
### AMPK Phosphorylation Proceeds Through Hierarchical Proteoform Cascades Revealed by Integrated Mass Spectrometry

Boris Krichel*[1,2,3,4,6], Hsin-Ju Chan[5], Liam Bandura[5], Zhan Gao[5], Man-Di Wang[5], Holden T. Rogers[5], Sean J. Mcilwain[7,8], Charlotte Uetrecht[4,6] and Ying Ge*[1,2,5]

[1] Department of Cell and Regenerative Biology, School of Medicine and Public Health, University of Wisconsin-Madison, Madison, Wisconsin, USA. [2]Human Proteomics Program, School of Medicine and Public Health, University of Wisconsin-Madison, Madison, Wisconsin, USA. [3] Faculty V: School of Life Sciences, University of Siegen, 57076 Siegen, Germany. [4]CSSB Centre for Structural Systems Biology, Deutsches Elektronen Synchroton DESY, Leibniz Institute of Virology and University of Lübeck, Notkestraße 85, 22607 Hamburg, Germany. [5]Department of Chemistry, University of Wisconsin-Madison, Madison, Wisconsin, USA. [6]Institute of Chemistry and Metabolomics. University of Lübeck, 23562 Lübeck, Germany. [7]Department of Biostatistics and Medical Informatics, University of Wisconsin-Madison, Madison, WI 53705, United States. [8]University of Wisconsin Carbone Cancer Center, University of Wisconsin-Madison, Madison, WI 53705, United States.

**Supplementary Table 1: Protein sequences of AMPK α1, β1 and γ1 subunits as used in the experiments in FASTA format.**

Sequences are identical to UniProt entries Q13131 AAPK1_HUMAN, Q9Y478 AAKB1_HUMAN, and P54619 AAKG1_HUMAN. In our recombinant protein, the first 10 amino acids were removed according to a naturally occurring AMPK variant and a His-6 tag with linker was added (blue). According to reviewed UniProt Q13131 AAPK1_HUMAN, residues were counted so the main activation site is T183. The targeted positions for site-directed mutations K56N and T183A highlighted in green. The N-terminal methionine (red) in α1 and β1 undergoes post-translational processing in *E. coli*.

| **AMPK protein subunit sequences in FASTA format** |
|---|
| >P7_AMPK_alpha1<br>MSYYHHHHHHEGVRTMATAEKQKHDGRVKIGHYILGDTLGVGTFGKVKVGKHELTGHKV<br>AVKILNRQKIRSLDVVGKIRREIQNLKLFRHPHIIKLYQVISTPSDIFMVMEYVSGGELFDYIC<br>KNGRLDEKESRRLFQQILSGVDYCHRHMVVHRDLKPENVLLDAHMNAKIADFGLSNMMS<br>DGEFLRTSCGSPNYAAPEVISGRLYAGPEVDIWSSGVILYALLCGTLPFDDDHVPTLFKKI<br>CDGIFYTPQYLNPSVISLLKHMLQVDPMKRATIKDIREHEWFKQDLPKYLFPEDPSYSSTM<br>IDDEALKEVCEKFECSEEEVLSCLYNRNHQDPLAVAYHLIIDNRRIMNEAKDFYLATSPPD<br>SFLDDHHLTRPHPERVPFLVAETPRARHTLDELNPQKSKHQGVRKAKWHLGIRSQSRPN<br>DIMAEVCRAIKQLDYEWKVVNPYYLRVRRKNPVTSTYSKMSLQLYQVDSRTYLLDFRSID<br>DEITEAKSGTATPQRSGSVSNYRSCQRSDSDAEAQGKSSEVSLTSSVTSLDSSPVDLTP<br>RPGSHTIEFFEMCANLIKILAQ* |
| >P7_AMPK_beta1<br>MGNTSSERAALERHGGHKTPRRDSSGGTKDGDRPKILMDSPEDADLFHSEEIKAPEKEE<br>FLAWQHDLEVNDKAPAQARPTVFRWTGGGKEVYLSGSFNNWSKLPLTRSHNNFVAILDL<br>PEGEHQYKFFVDGQWTHDPSEPIVTSQLGTVNNIIQVKKTDFEVFDALMVDSQKCSDVSE<br>LSSSPPGPYHQEPYVCKPEERFRAPPILPPHLLQVILNKDTGISCDPALLPEPNHVMLNHL<br>YALSIKDGVMVLSATHRYKKKYVTTLLYKPI |
| >P7_AMPK_gamma1<br>METVISSDSSPAVENEHPQETPESNNSVYTSFMKSHRCYDLIPTSSKLVVFDTSLQVKKA<br>FFALVTNGVRAAPLWDSKKQSFVGMLTITDFINILHRYYKSALVQIYELEEHKIETWREVYL<br>QDSFKPLVCISPNASLFDAVSSLIRNKIHRLPVIDPESGNTLYILTHKRILKFLKLFITEFPKPE<br>FMSKSLEELQIGTYANIAMVRTTTPVYVALGIFVQHRVSALPVVDEKGRVVDIYSKFDVINL<br>AAEKTYNNLDVSVTKALQHRSHYFEGVLKCYLHETLETIINRLVEAEVHRLVVVDENDVVK<br>GIVSLSDILQALVLTGGEKKP* |

**Supplementary Table 2: Intact masses of AMPK subunits measured by RPLC-MS.**

Values represent the most abundant masses of unmodified protein subunits at 0 min time point. Data were acquired in triplicates using the Impact II Q-TOF mass spectrometer. Experimental (Expt'l) masses were determined using the sum peak algorithm to find the most abundant mass. Calculated (Calc'd) masses represent the most abundant theoretical mass determined with ProtPi. SD = Standard Deviation (n=3).

| AMPK Subunit | Expt’l Replicate 1 (Da) | Expt’l Replicate 2 (Da) | Expt’l Replicate 3 (Da) | Expt’l Mean (Da)± SD (Da) | Calc’d (Da) |
|---|---|---|---|---|---|
| α1 | 64587.6 | 64584.9 | 64586.6 | 64586.4 ± 1.4 | 64586.0 |
| β1 | 30250.12 | 30250.12 | 30250.13 | 30250.12 ± 0.01 | 30250.28 |
| γ1 | 37578.59 | 37578.60 | 37578.66 | 37578.62 ± 0.04 | 37578.83 |

**Supplementary Table 3: Phosphopeptide quality parameters from MS-Fragger raw data search results.**

Listed are the phosphopeptides used for kinetic growth curves. The table shows the residue with the most likely modification (Res.) and their mass increase in Dalton (Mod.), the observed m/z value (*m/z*), the expectation value (Exp.), which is considered excellent when <0.01. Also included are the Peptide Prophet Probability (PPP) from 0 to 1, and the STY localization value indicating the probability (0 to 1) of the marked site being modified (Loc.). Phosphopeptide pairs were selected based on spectral quality, site localization confidence, and consistent inverse correlation behavior.

| Res. | Mod. | *m/z* | Exp. | PPP | Loc. |
|---|---|---|---|---|---|
| α1 S136 | LFQQILS[+80]GVDYC[+57]HR | 455.0 | 1.3E-01 | 0.83 | 0.67 |
| α1 S176 | IADFGLSNMMS[+80]DGEFLR | 991.9 | 1.4E-09 | 1.00 | 0.95 |
| α1 T183 | T[+80]SC[+57]GSPNYAAPEVISGR | 923.4 | 3.2E-11 | 1.00 | 0.94 |
| α1 S251 | IC[+57]DGIFYTPQYLNPS[+80]VISLLK | 1261.1 | 6.4E-09 | 1.00 | 0.95 |
| α1 T355 | DFYLAT[+80]SPPDSFLDDHHLTRPHPER | 761.6 | 1.4E-07 | 1.00 | 0.95 |
| α1 S356 | DFYLATS[+80]PPDSFLDDHHLTRPHPER | 609.5 | 5.9E-06 | 1.00 | 0.51 |
| α1 S360 | DFYLATSPPDS[+80]FLDDHHLTRPHPER | 761.6 | 9.4E-10 | 1.00 | 0.93 |
| α1 T388 | HT[+80]LDELNPQK | 637.8 | 9.9E-08 | 1.00 | 1.00 |
| α1 S397 | HTLDELNPQKS[+80]K | 745.4 | 1.9E-05 | 1.00 | 0.95 |
| α1 S496 | SGS[+80]VSNYR | 948.4 | 1.3E-03 | 0.99 | 0.91 |
| α1 S516 | S[+80]SEVSLTSSVTSLDSSPVDLTPR | 815.4 | 9.0E-04 | 1.00 | 0.41 |
| α1 S524 | SSEVSLTSS[+80]VTSLDSSPVDLTPR | 1223.1 | 1.6E-12 | 1.00 | 0.94 |
| α1 T526 | SSEVSLTSSVT[+80]SLDSSPVDLTPR | 1222.6 | 1.0E-09 | 1.00 | 0.94 |
| β1 S96 | EVYLSGS[+80]FNNWSK | 805.8 | 1.6E-06 | 1.00 | 0.94 |
| β1 S101 | EVYLSGSFNNWS[+80]KLPLTR | 1096.0 | 2.3E-07 | 1.00 | 0.94 |
| β1 S108 | S[+80]HNNFVAILDLPEGEHQYK | 764.4 | 1.0E-14 | 1.00 | 0.94 |
| β1 T148 | FFVDGQWTHDPSEPIVTSQLGT[+80]VN NIIQVK | 1150.6 | 6.3E-10 | 1.00 | 0.95 |
| γ1 S9 | METVISSDS[+80]SPAVENEHPQETPESN NSVYTSFM[+16]K | 1289.9 | 2.0E-14 | 1.00 | 0.94 |

**Supplementary Table 4: Consistent retention times of different phosphopeptide pairs used for quantification.**

Listed are the phosphopeptides used for kinetic growth curves. Data from Skyline extracted ion chromatograms at the 50 min timepoint (n=3). Shown are the residues with the most likely modification (Res.), the modified peptide sequence and their mass increase in Dalton (Mod.), the average retention time in minutes for the parent peptide (x̄ Pep RT), the average retention time in minutes for the phosphorylated peptide (x̄ pPep RT), and the retention time difference in minutes between parent and phosphorylated forms (ΔRT). Elution times were highly reproducible between replicates (±0.1 min within triplicates).

| Res. | Mod. | x̄ Pep RT (min) | x̄ pPep. RT (min) | Δ RT (min) |
|---|---|---|---|---|
| α1 S136 | R.LFQQILS[+80]GVDYC[+57]HR.H [135, 148] | 41.6 | 43.5 | -1.9 |
| α1 S176 | IADFGLSNMMS[+80]DGEFLR | 51.6 | 52.2 | -0.6 |
| α1 T183 | T[+80]SC[+57]GSPNYAAPEVISGR | 29.8 | 31.9 | -2.1 |
| α1 S251 | IC[+80]DGIFYTPQYLNPS[+80]VISLLK | 53.7 | 54.1 | -0.4 |
| α1 T355, S356, S360 | DFYLAT[+80]SPPDSFLDDHHLTRPHPER | 39.9 | 41.5 | -1.6 |
| α1 T388 | HT[+80]LDELNPQK | 19.8 | 21.3 | -1.5 |
| α1 S397 | HTLDELNPQKS[+80]K | 17.5 | 18.9 | -1.4 |
| α1 S496 | SG[+80]SVSNYR | 11.1 | 12.2 | -1.1 |
| α1 S516 | S[+80]SEVSLTSSVTSLDSSPVDLTPR | 45.9 | 47.9 | -2.0 |
| α1 S524 | SSEVSLTSS[+80]VTSLDSSPVDLTPR | 45.9 | 45.3 | 0.6 |
| α1 T526 | SSEVSLTSSVT[+80]SLDSSPVDLTPR | 45.9 | 46.7 | -0.8 |
| β1 S96 | EVYLSGS[+80]FNNWSK | 38.6 | 40.3 | -1.7 |
| β1 S101 | EVYLSGSFNNWS[+80]KLPLTR | 47.2 | 50.0 | -2.8 |
| β1 S108 | S[+80]HNNFVAILDLPEGEHQYK | 41.1 | 44.6 | -3.5 |
| β1 T148 | FFVDGQWTHDPSEPIVTSQLGT[+80]VNNIIQVK | 51.3 | 51.3 | -0.1 |
| γ1 S9 | METVISSDS[+80]SPAVENEHPQETPESNNSVY TSFM[+16]K | 40.1 | 41.7 | -1.6 |

**Supplementary Table 5: Analysis of unphosphorylated and phosphorylated peptide pairs and their kinetic curves.**

Listed are the phosphorylation site position, extent of phosphorylation (fraction of phosphorylated peptide calculated from maximum to minimum normalized AUC values), rate constants (k) from kinetic model fitting, and $R^2$ values indicating goodness of fit. Unphosphorylated peptides were fit with one-phase decay models and phosphorylated peptides with one-phase association models. See Methods for detailed analysis procedures.

| Position | Extent | k unphos. fit [$min^{-1}$] | $R^2$ unphos. fit | k phos. fit. [$min^{-1}$] | $R^2$ phos. fit |
|---|---|---|---|---|---|
| α S136 | 0.09 | 0.071 ± 0.15 | 0.788 | 0.005 ± 0.002 | 0.998 |
| α S176 | 0.624 | 0.053 ± 0.014 | 0.994 | 0.067 ± 0.09 | 0.893 |
| α T183 | 0.842 | 0.077 ± 0.005 | 0.998 | 0.110 ± 0.09 | 0.957 |
| α S251 | 0.104 | 0.036 ± 0.15 | 0.648 | 0.015 ± 0.15 | 0.818 |
| α T355/S356/S360 | 0.23 | 1.13 ± 0.56 | 0.762 | 0.005 ± 0.005 | 0.99 |
| α T388 | 0.346 | 0.007 ± 0.004 | 0.957 | 0.017 ± 0.007 | 0.992 |
| α S397 | 0.347 | 0.002 ± 0.04 | 0.648 | 0.002 ± 0.04 | 0.969 |
| α S496 | 0.993 | 0.161 ± 0.046 | 0.991 | 0.0001 ± 0.002 | 0.878 |
| α S516 | 0.812 | 0.008 ± 0.008 | 0.971 | 0.115 ± 0.14 | 0.908 |
| α S524 | 0.812 | 0.008 ± 0.008 | 0.971 | 0.084 ± 0.05 | 0.967 |
| α T526 | 0.812 | 0.008 ± 0.008 | 0.971 | 0.225 ± 0.11 | 0.584 |
| β S96 | 0.293 | 0.017 ± 0.04 | 0.857 | 0.020 ± 0.02 | 0.984 |
| β S101 | 0.292 | 0.226 ± 0.72 | 0.714 | 0.012 ± 0.008 | 0.995 |
| β S108 | 0.981 | 0.051 ± 0.038 | 0.983 | 0.062 ± 0.04 | 0.971 |
| β T148 | 0.45 | 0.077 ± 0.083 | 0.937 | 0.018 ± 0.02 | 0.951 |
| γ S9 | 0.803 | 0.055 ± 0.027 | 0.986 | 0.021 ± 0.009 | 0.984 |

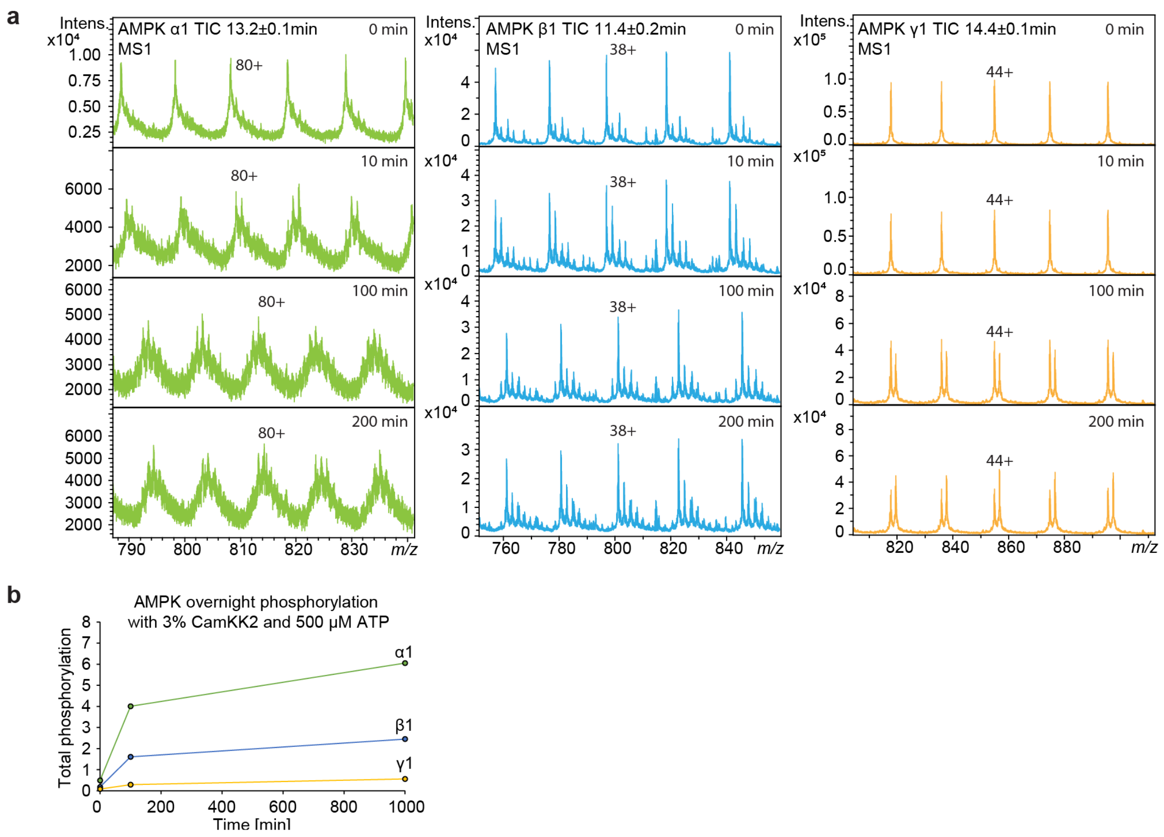

**Supplementary Figure 1: Extended intact protein AMPK proteoform analysis**.

**(a)** Representative raw mass spectra showing temporal progression of AMPK subunit phosphorylation by CamKK2 at four timepoints (0, 10, 100, and 200 min) with the five most intense charge states shown and one representative charge state labeled (left: α1, middle: β1, right: γ1). Total ion chromatogram (TIC) integration windows shown for spectral summary. **(b)** Extended phosphorylation time course at 0, 100, and 1000 min. The modest increase between 100 and 1000 min validates that the 200 min time point captured the majority of phosphorylation events.

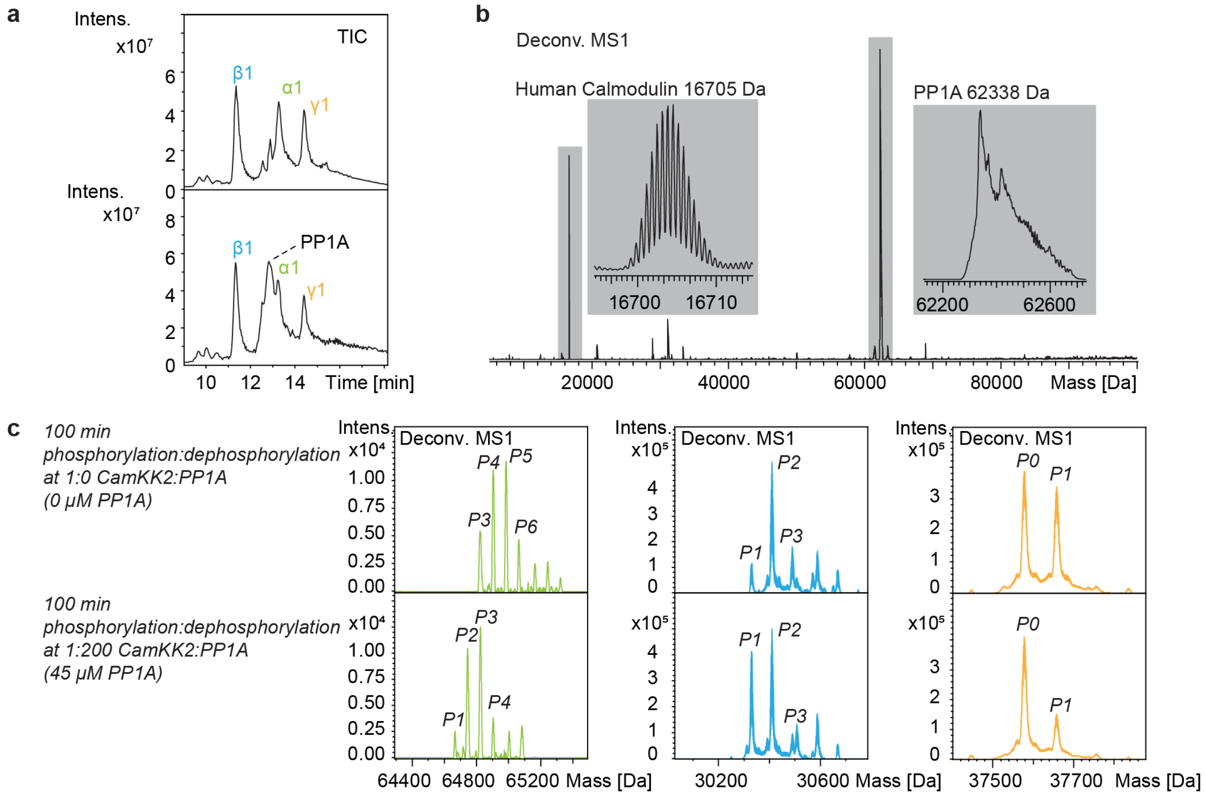

**Supplementary Figure 2: PP1A phosphatase-mediated dephosphorylation characterization.**

**(a)** Total ion chromatograms comparing AMPK samples without and with 45 µM PP1A, showing distinct phosphatase peak appearance. **(b)** MS1 shows coeluting human calmodulin and PP1A. **(c)** Representative deconvoluted mass spectra comparing 100 min phosphorylation products without (top) and with PP1A treatment (bottom) for all three subunits (left: α1, middle: β1, right: γ1), with phosphorylation states numerically labeled.

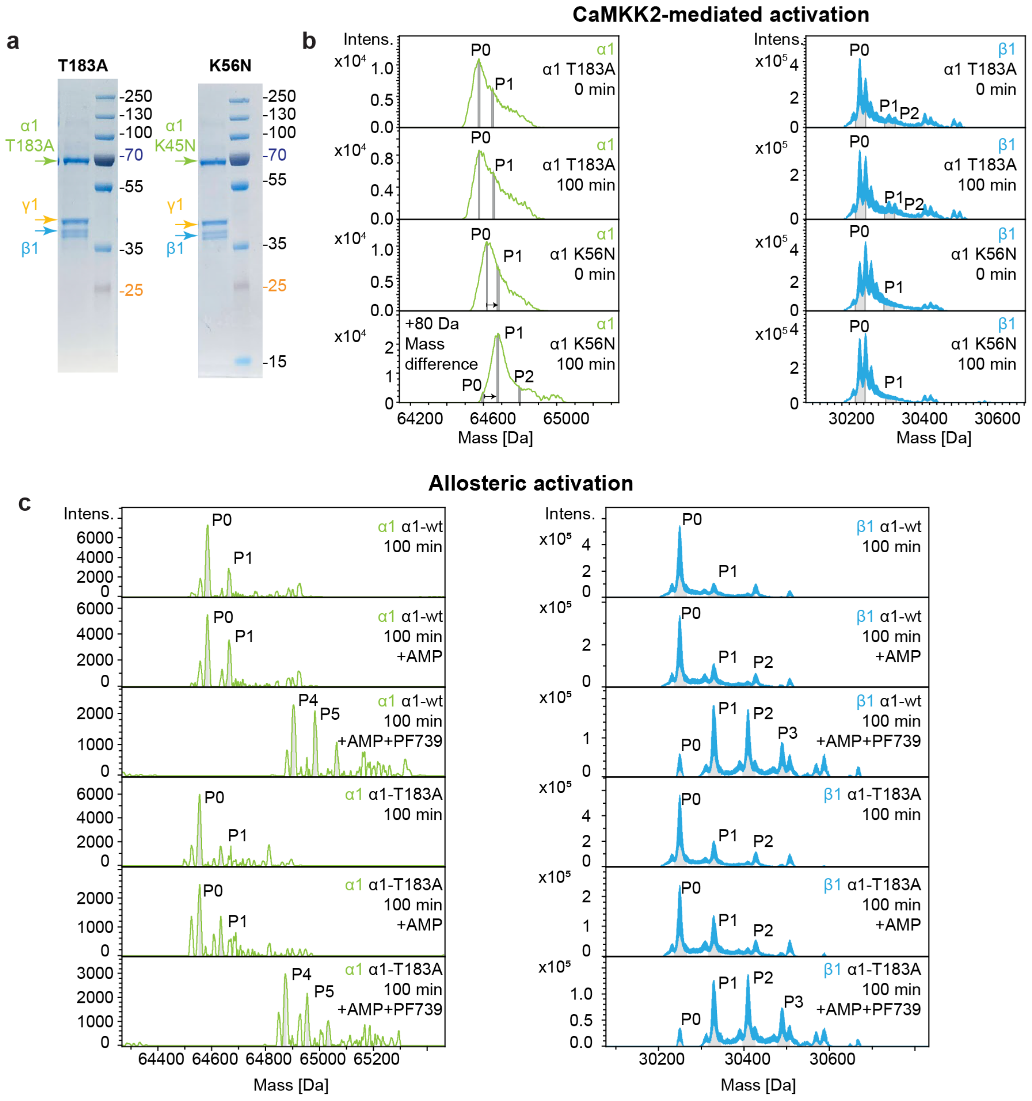

**d**

| | α1 wt P0 | α1 T183A P0 | β1 P0 (from wt) | β1 P0 (from α1 T183A) |
|---|---|---|---|---|
| Calc'd | 64586.0 Da | 64556.6 Da | 30250.3 Da | 30250.3 Da |
| Expt'l | 64586.4 Da | 64556.4 Da | 30250.1 Da | 30250.1 Da |
| Δ | 0.4 Da | 0.2 Da | 0.2 Da | 0.2 Da |

**Supplementary Figure 3: Mutant AMPK proteoform analysis reveals distinct phosphorylation mechanisms.**

**(a)** SDS-PAGE analysis of IMAC-purified T183A and K56N mutants shows purification to homogeneity. **(b)** Deconvoluted mass spectra of α1 (left) and β1 (right) subunits comparing T183A and K56N mutants under CaMKK2 activation conditions (0 and 100 min). AMPK β1 subunit reveals detailed oxidation states (+16 Da), while α1 subunit oxidation is indicated by mass differences averaging one oxidation for T183A and three oxidations for K56N. K56N mutant shows CaMKK2-mediated α1 phosphorylation at 100 min despite catalytic inactivity, whereas T183A lacks activation loop phosphorylation. Limited spectral resolution for α1 subunit required peak shoulder analysis based on expected Δm of phosphorylation states. Despite oxidation, the main result is clearly evident - the mass shift of approx. 80 Da indicated by the arrow arising from one phosphorylation in α1 K56N. **(c)** Comparative mass spectra of wild-type and T183A mutants under allosteric activation conditions showing α1 (left) and β1 (right) subunit phosphorylation. Conditions include: 100 min incubation, AMP addition, and combined AMP plus PF-739 treatment. Mass differences (0.4-0.7 Da) prove that no oxidation is present in this protein preparation. PF-739 allosteric activation induces extensive autophosphorylation comparable to CaMKK2-mediated activation, independent of activation loop phosphorylation status. **(d)** Determined masses confirming wildtype and mutated (T183A) protein sequence. Experimental (Expt'l) masses determined by manual isotopologue picking and calculated (Calc'd) masses determined with ProtParam.

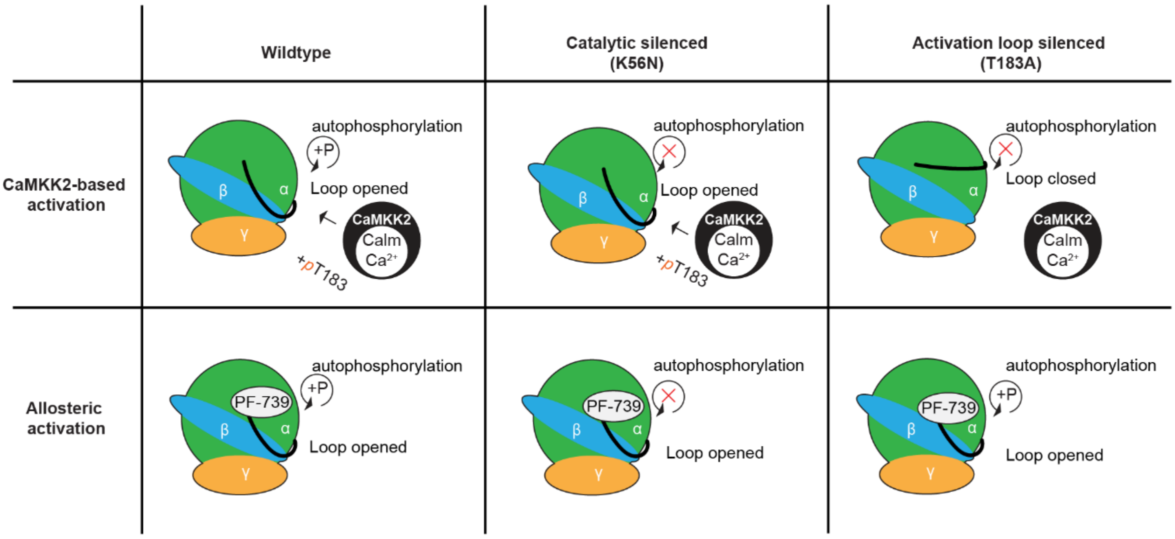

**Supplementary Figure 4: Schematic representation of AMPK activation mechanisms.**

Schematic representation of AMPK activation mechanisms comparing wild-type, catalytically inactive (K56N), and activation loop-deficient (T183A) mutants. Upper panel illustrates CaMKK2-mediated activation showing differential activation loop phosphorylation and autophosphorylation capacity. Lower panel depicts allosteric activation pathway demonstrating PF-739-induced conformational changes that bypass canonical phosphorylation requirements while maintaining autophosphorylation activity across all variants.

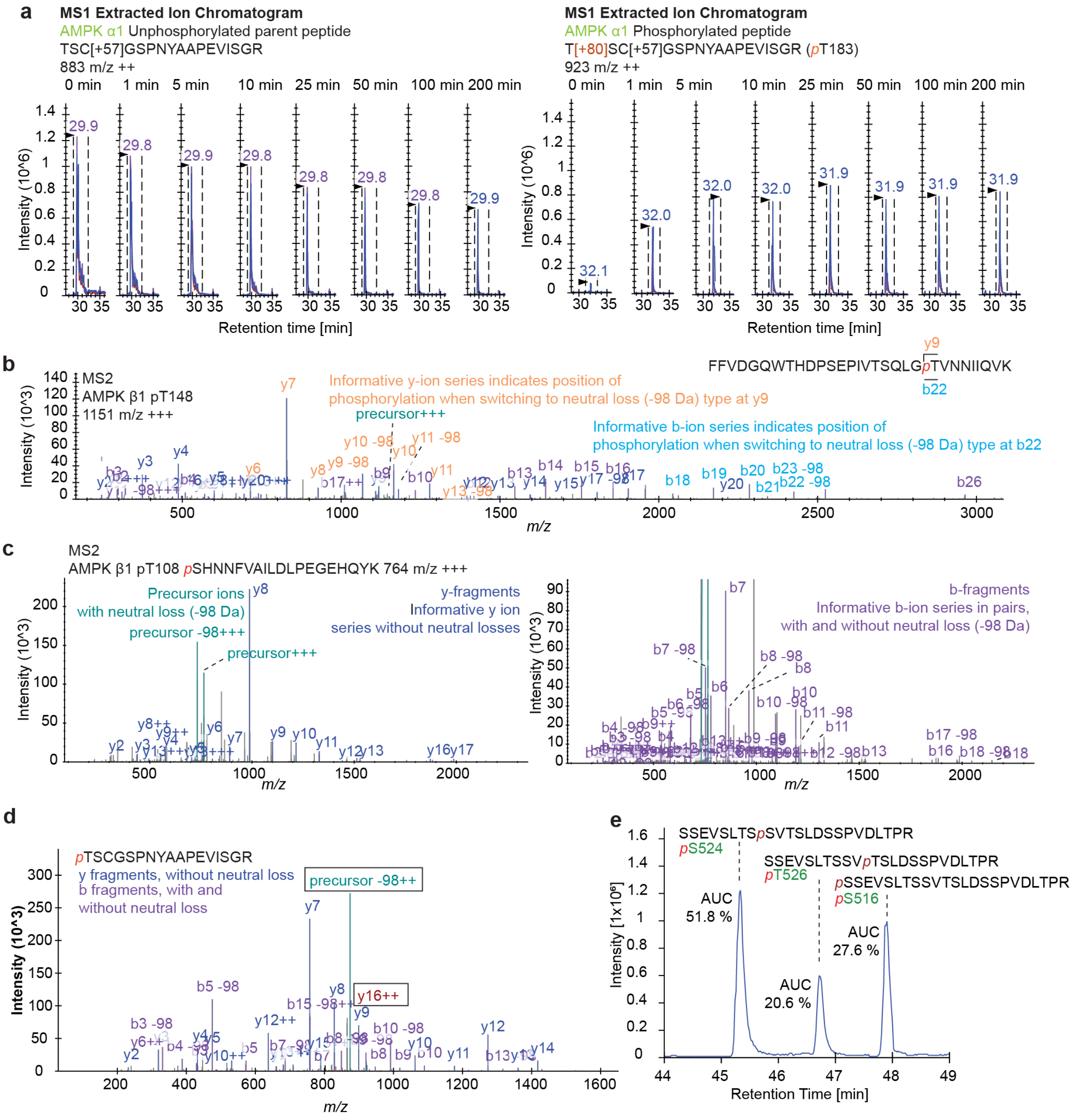

**Supplementary Figure 5: Representative data illustrating the bottom-up workflow for site-specific phosphorylation kinetics.**

**(a)** Extracted ion chromatograms from a single replicate across eight sampling timepoints (0-200 min) showing parent peptide and phosphorylated peptide corresponding to α1-T183 phosphorylation. *m/z* values and retention times labeled. **(b)** MS2 library match for phosphorylation at β1 T158. The fragment pattern and ion series switch to neutral loss masses unequivocally localizes the phosphorylation site. **(c)** MS2 library match of N-terminal modification at β1 S108. Highlighted are y-fragments (left) and b-fragments (right) that indicate the MS2 library match. **(d)** MS2 library match of phosphorylation at α1-T183. While N-terminal phosphorylation is indicated by b- and y-fragment pattern and neutral loss distribution, the exact location is only indicated by a low-intensity y16 fragment and its absence of neutral loss. However, since this represents the main AMPK activation site, this location has been confidently and repeatedly assigned through complementary methods. **(e)** A 23-residue phosphopeptide containing S516, S524, and T526 separates into three distinct chromatographic peaks. Peak area quantification (shown as % AUC) demonstrates site-specific resolution of closely spaced phosphorylation sites.

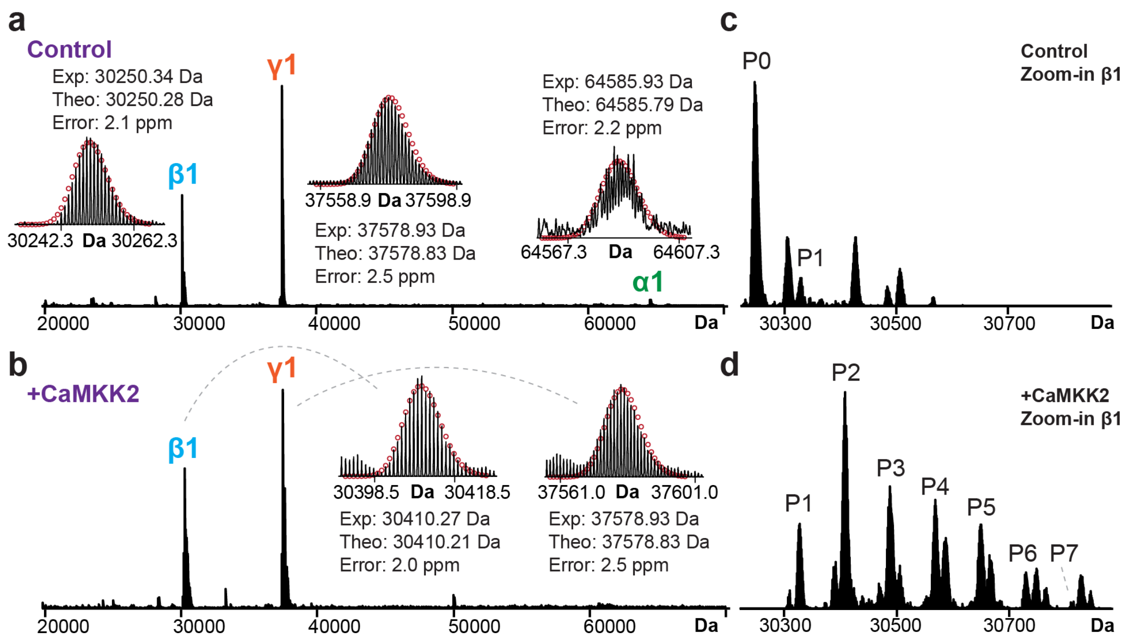

**Supplementary Figure 6: Ultra high-resolution top-down mass spectrometry of AMPK subunits acquired on FTICR via direct infusion.**

Deconvoluted mass spectra of **(a)** untreated control and **(b)** CaMKK2-treated AMPK. Experimental masses, theoretical masses, and mass errors of the most abundant proteoforms for each subunit are reported. Isotopic fittings are shown in red circles. Zoomed-in views of the AMPK β1 from **(c)** untreated control and **(d)** CaMKK2-treated AMPK. Phosphoproteoforms are labeled as P0-P7 according to phosphorylation numbers.

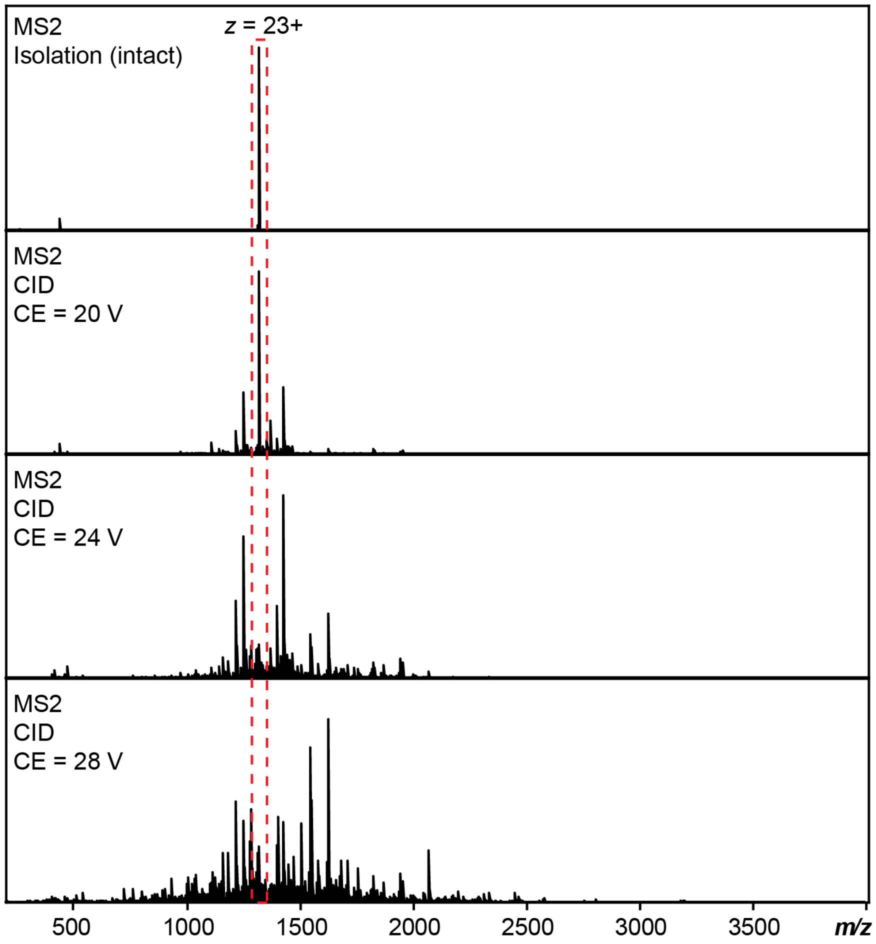

**Supplementary Figure 7: MS/MS characterization of the bis-phosphorylated AMPK β1 subunit using collision-induced dissociation (CID).**

The bis-phosphorylated AMPK β1 (z = 23+) was isolated, and a CID energy of 20 V, 24 V, and 28 V were applied in the collision cell, respectively.

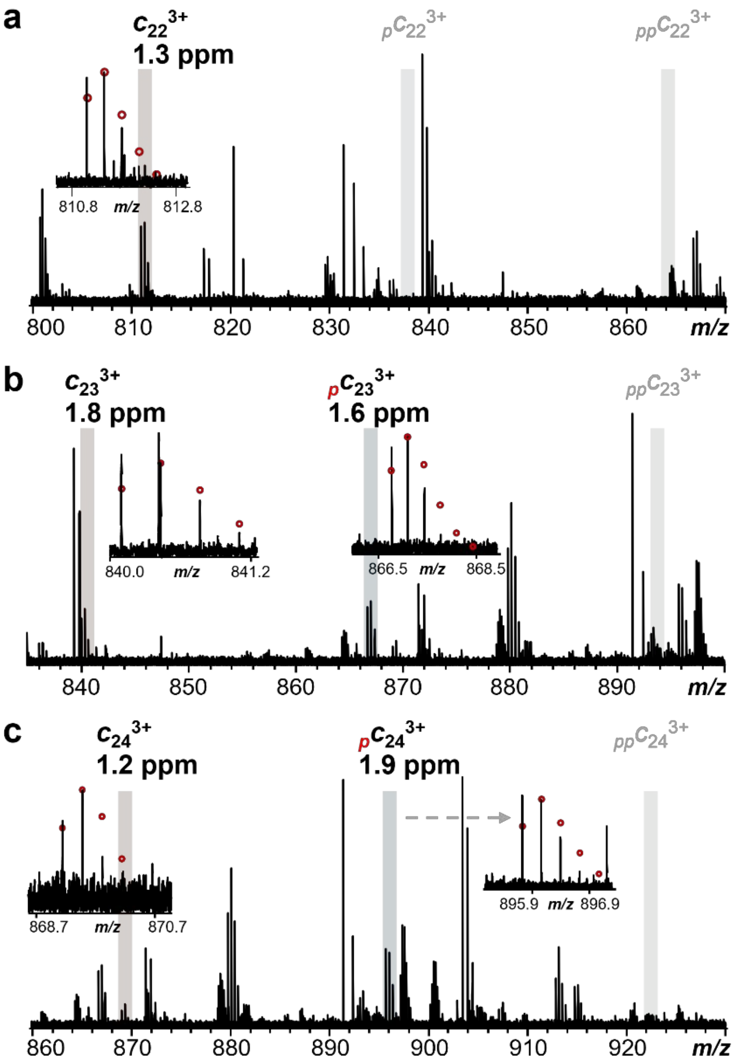

**d**

| Ion | Charge | Intensity | | | %P Total | %P Site |
|---|---|---|---|---|---|---|
| | | Un | Mono | Bis | | |
| c22 | 3+ | 1.5E+07 | 0.0E+00 | 0.0E+00 | 0 | 0 |
| c23 | 3+ | 9.0E+06 | 1.3E+07 | 0.0E+00 | 60 | 60 |
| c24 | 3+ | 5.2E+06 | 1.6E+07 | 0.0E+00 | 76 | 16 |

**Supplementary Figure 8: Localization of phosphorylation sites on bis-phosphorylated AMPK β1 by electron-capture dissociation (ECD).**

Representative ECD MS/MS fragment spectra of (a) $c_{22}^{3+}$, **(b)** $c_{23}^{3+}$, **(c)** $c_{24}^{3+}$. The $_{p}$c and $_{pp}$c represent mono- and bis-phosphorylated c ions, respectively. Shift from fragment pattern $c_{22}$, $c_{23}$, $c_{24}$ versus assigned positions pS24, pS25 due to N-terminal glycine cleavage. Isotopomer peak profiles of the identified fragment ions are expanded in each spectrum. Circles represent the theoretical isotopic distribution corresponding to the assigned molecular weight. The mass accuracy for each identified fragment is labeled in ppm. **(d)** Intensity of un-/mono-/bis-phosphorylated fragments. The percentage of phosphorylated fragments for each c ion group (%P total) was calculated, and the phosphorylation percentage at each amino acid residue (%P site) was derived from the %P total.
